# A developmentally scheduled inhibitory polarity circuit centers the stomatal pore

**DOI:** 10.64898/2026.10.31.763351

**Authors:** Alexis Lebecq, Eva-Sophie Wallner, Dominique C. Bergmann

## Abstract

Symmetry in living systems often arises from coordinated asymmetries at smaller scales. The guard cells of stomata--bilaterally symmetric epidermal valves that mediate plant-atmosphere gas exchange- -show coordinated asymmetries in pore placement and wall modification. Stomatal form and function are deeply conserved, yet molecular mechanisms defining coordinated cellular asymmetry are unknown. Here we show that pore position arises from a combination of pole avoidance and gradient centering in the membranes and wall dividing sister guard cells. Boundaries are established by spatiotemporal restriction of the receptor-like kinase MUSTACHES (MUS), which directs the disordered polarity protein OCTOPUS-LIKE2 (OPL2) to pore-forming regions. OPL2 peaks correspond to a focus of microtubules, and, from this position, the pore emerges. Experimental rewiring of the MUS-OPL2 polarity module profoundly impacts stomatal morphogenesis, eliminating, shifting, or even redirecting the pore to the opposite wall. These findings reveal a stomata-specific integration and elaboration of core polarity and centering programs, and challenge previous models of stomatal morphogenesis.

## INTRODUCTION

Symmetry is a recurring organizational feature of living systems, yet it often arises from coordinated asymmetry rather than uniformity. In multicellular organisms, polarity circuits partition individual cells into distinct biochemical domains; when sister cells align these domains across a shared interface, bilateral symmetry emerges at tissue or organ scales (1, 2). Stomata, the epidermal valves that mediate plant-atmosphere gas exchange, present a particularly attractive system for studying coupled asymmetries. Stomata feature a pair of guard cells (GCs) surrounding a central aperture. Already in 450-million-year-old fossils, the GCs appear as mirror-symmetric, kidney-shaped cells (3), and this morphology remains widespread in extant plants, including Arabidopsis (4). GCs are sister cells, born from the division of a guard mother cell (GMC) (5). Each GC undergoes polarized morphogenesis while maintaining mirror-image coordination — depositing specialized wall material and expanding through turgor pressure to produce a pore centered on the shared ‘ventral’ or ‘inner’ wall (Fig 1A 6). Although GC wall composition and mechanics have been studied extensively (6-8), the mechanisms that coordinate two sister cells to generate the iconic stomatal geometry are not known.

**Figure 1:**
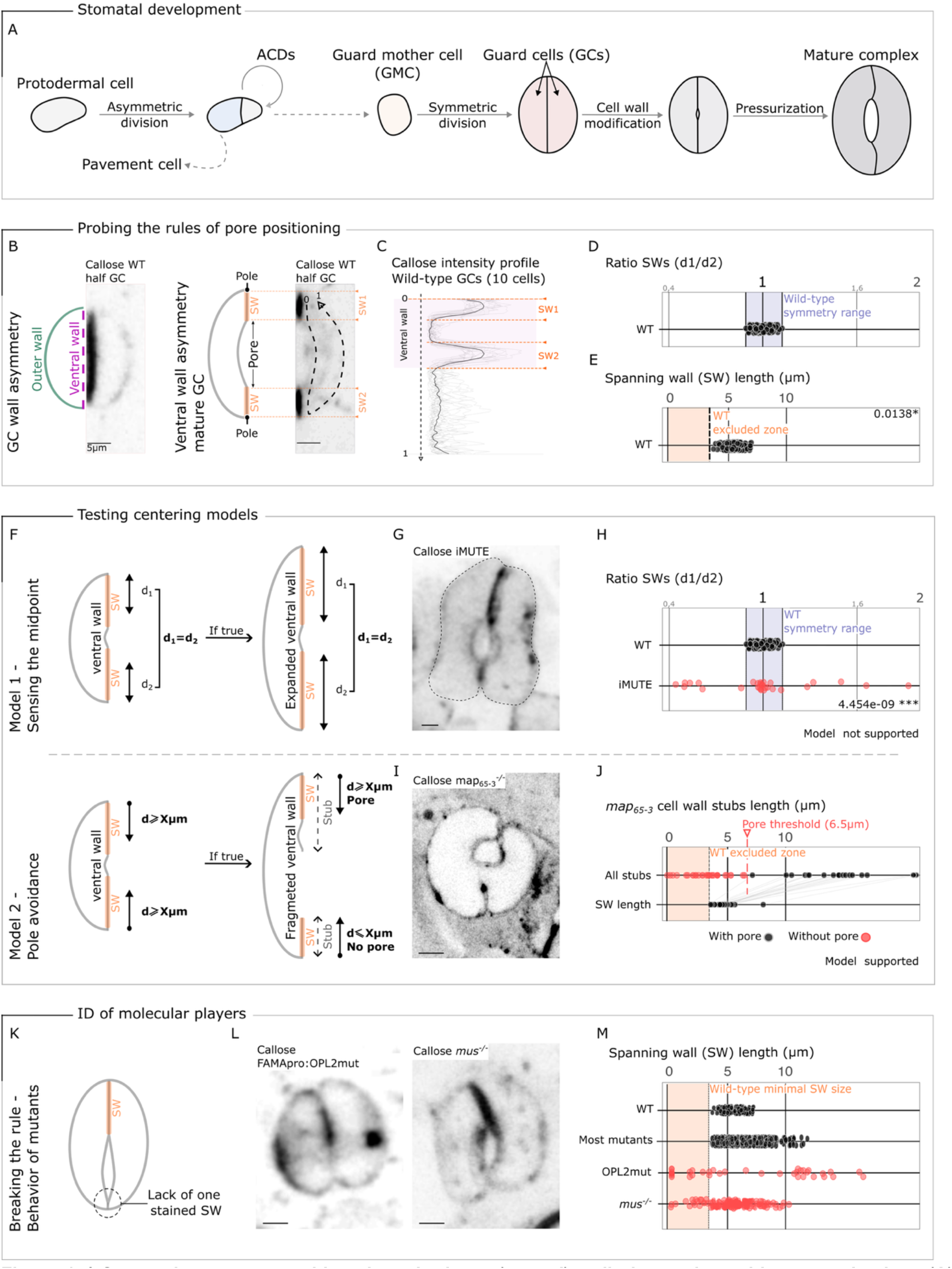
Stomatal pores are positioned on the inner (ventral) wall via a pole avoidance mechanism. **(A)** Schematic of the stomatal development pathway, highlighting Guard Mother Cell (GMC) specification, symmetric GMC division into guard cells (GCs), and subsequent cell wall remodeling and turgor-driven maturation. **(B)** Aniline blue staining of callose preferentially marks the ventral (magenta) GC walls established at GMC division, distinguishing them from the outer walls (green). Orange dotted lines demarcate the two spanning walls (SW1, SW2) flanking the pore; Arrow indicates the orientation of the intensity profile used in (C). **(C)** Averaged callose intensity profiles across the ventral wall (n = 10 GCs). **(D)** SW length symmetry (d1/d2) from 7-dpg wild-type cotyledons (n = 117 GCs, 10 cotyledons); blue shading, observed symmetry range. **(E)** Distribution of individual SW lengths from 7-dpg wild-type cotyledons (n = 234 GCs); revealing a minimal SW size (orange shading here and in J, M marks zone excluded in WT) * Represents significance, bootstrap-based one-sample probability test. **(F)** Midpoint sensing model: prediction if pores are positioned by equalizing SW distances (d1= d2). Pole avoidance model: prediction if pore position requires SW length X m. **(G)** Representative iMUTE-induced stoma∼1.5× larger than WT at 3dpg. **(H)** SW symmetry in wild-type (n = 117 GCs) and iMUTE-induced stomata (n = 30 GCs, 5 cotyledons, red). **(I)** Representative *map65-3* stoma at 7dpg showing stubs that can or cannot form a pore. **(J)** Wall stub lengths in *map65-3* stratified by pore presence (n = 18 GCs, 13 cotyledons) or absence (n = 29, 17 cotyledons, red), with SW lengths for pore-forming stubs **(K)** Expected phenotypes when pore-positioning rules are violated. **(L)** Representative *FAMApro:OPL2mut* and *mustaches* (*mus*) stomata. **(M)** SW length distributions in 7-dpg wild-type (n = 234), mutants in other candidate genes (n = 752, 16 genotypes), *FAMApro:OPL2mut* (n = 42, from 5 cotyledons), and *mus* (n = 206, from 10 cotyledons); red dots indicate stomata with mispositioned pores. Scale bars, 5m for all images. *Associated supplemental figures: S1-S5*.

Here we combine genetics, timelapse imaging, and synthetic rewiring of cell polarity to show that stomatal pore centering involves pole avoidance and gradient formation at the interface of developing GCs. The molecular components of this strategy are the transmembrane receptor-like kinase MUSTACHES (MUS) and the intrinsically disordered scaffold protein OCTOPUS-LIKE 2 (OPL2). Cell-cycle-restricted expression of MUS enriches it at GC poles, where it generates a graded, diffusible inhibitory field that prevents OPL2 membrane accumulation. Within a permitted zone, OPL2 assembles in a gradient whose peak marks a focus of cortical microtubules and site of pore initiation. This MUS-OPL2-MT circuit is sufficient to organize morphogenesis and additional molecular asymmetries to generate the mirror-symmetric kidney-shaped GCs that comprise the functional stoma.

## RESULTS

Stomata are historically described as having centered pores (9). If we are to discover the mechanisms underlying stomatal form, we need a more quantitative assessment of this claim; are pores truly centered, and are they centered initially, or does centering emerge during stomatal maturation? We took advantage of the fact that callose, easily detected by calcofluor staining, is enriched in the two ‘spanning walls’ (SWs) adjacent to the stomatal pore on the ventral face of mature GCs (Fig.1B, C). Pore position can be described by the ratio of SW lengths; if SW lengths are symmetric (d1/d2 = 1) the pore is centered. SW lengths were symmetric in young and in mature GCs of wild type Arabidopsis cotyledons (7 days post-germination, dpg) indicating that pores initiate and remain centered during stomatal maturation, despite substantial increases in ventral wall and pore length (Fig. 1D, fig. S1A–H). This organization was conserved across all 11 angiosperm species examined (fig. S2A). Because we measured absolute SW lengths, we also noticed that SWs and pore size were anticorrelated: following pore initiation, SW lengths decreased from ∼7µm to a minimum of ∼3µm as pores occupied an increasingly larger fraction of the ventral wall (Fig. 1E, fig. S1C– D). Decrease of SW length followed by persistence of this minimum suggested that pore positioning may result from inhibition at the ventral–outer wall junction. Such a ‘pole avoidance’ mechanism can be distinguished from direct sensing of the ventral wall midpoint by altering GC sizes, as midpoint-sensing should center the pore, regardless of ventral wall length (Fig. 1F, fig. S2B). When we generated ∼1.5-fold larger GCs via ectopic MUTE expression (10,11) (Fig. 1G, fig. S3), pore size remained largely unchanged, but pores were significantly less centered (Fig. 1H, fig. S3). In contrast, ‘pole avoidance’ predicts both reduced centering in elongated guard cells and a minimum wall length required for pore formation. Consistent with this, *map65-3* mutants, which produce discontinuous ventral wall stubs of variable length during GMC division (Fig. 1J; 12,13), formed pores only on stubs longer than ∼7 µm, leaving a minimum SW of ∼3 µm (Fig. 1K, fig. S4A). Together, these perturbations support pole avoidance as the mechanism underlying pore positioning and suggest that the same spatial constraint governs both pore initiation and pore expansion. What remains unresolved is the identity of the avoidance cue.

To identify the molecular basis of pole-avoiding behavior, we screened candidate mutants disrupting wall composition, cytoskeletal dynamics, vesicle trafficking, or cell polarity for violations of the SW size minimum. Pole-avoidance was robust to nearly all perturbations—including loss of *KATANIN, FERONIA*, and *BASL*— even when GC morphology was visibly altered in these mutants (fig. S4B-D). Two polarity-class mutants, however, broke the ∼3 µm minimum SW threshold: loss of the leucine-rich repeat receptor-like kinase MUSTACHES (MUS) (14), and a stabilized form of the intrinsically disordered polarity protein OCTOPUS-LIKE2 (OPL2mut) (15, 16). In *mus* mutants, the pore was shifted toward one of the poles. OPL2mut stomata largely failed to form a pore, but ectopic splits appeared near the poles of two older GCs (Fig. 1L-N, fig. S4, S5). OPL2 and MUS are therefore candidate components of the pole-avoidance machinery.

Previous work showed that MUS and OPL2 occupy polar positions in dividing GMCs and GCs (14, 16) and using new double-reporter lines, we confirmed their opposing outer-inner polarities (Fig. 2A). A conceptually straightforward (though technically challenging) test of the importance of MUS and OPL2’s opposing polar localization for stomatal morphogenesis would be to eliminate or swap their positions. Two main strategies have been used to relocalize polar proteins in plants: domain swaps between oppositely localized proteins (e.g.17) and manipulation of the cellular processes used to direct and maintain proteins at discrete locations (e.g. vesicle trafficking, the cytoskeleton, or the cell wall (18)). While pursuing these strategies with MUS and OPL2, we serendipitously found that simply changing the cell type and cell-cycle timing of MUS expression was sufficient to alter its localization. When expressed under different promoters active at successive stages during GMC to GC differentiation, the transmembrane RLK protein MUS could be directed to the plasma membrane adjacent to the outer walls (wild-type polarity), ventral walls (inverted polarity), or everywhere (Fig. 2B-D, fig. S6). This behavior suggests that MUS localization emerges from developmental scheduling coupled to bulk secretory flow rather than intrinsic polar sorting motifs. MUS can be directed to OPL2’s native (ventral) domain because transmembrane proteins are preferentially trafficked to the cell plate forming the nascent wall and membranes during cytokinesis (19).

**Figure 2:**
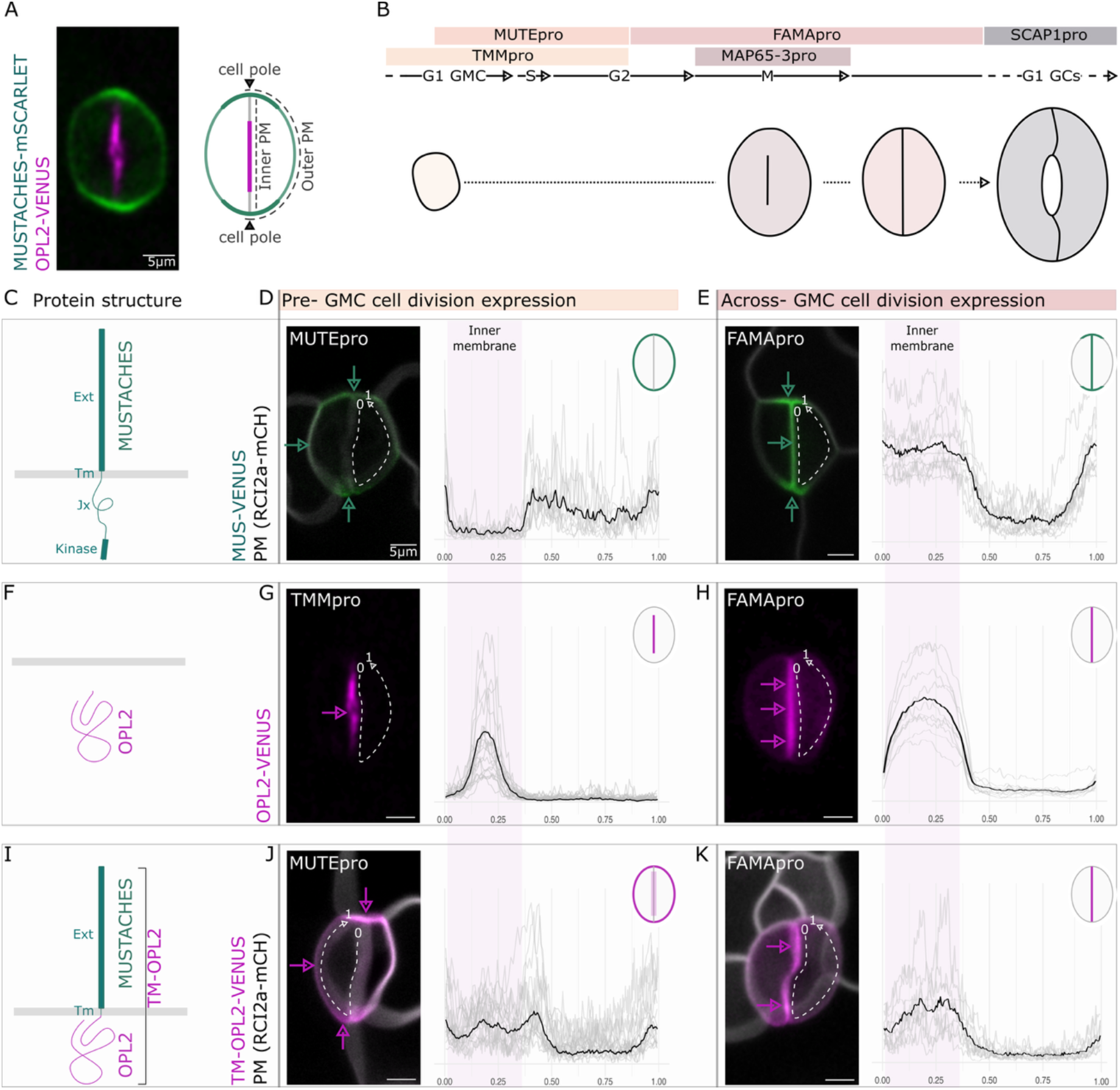
Polar localization of MUS and OPL2 can be rewired by altering expression timing and membrane attachment. **(A)** Cartoon and representative images of MUS (green) and OPL2 (magenta) in wild-type guard cells. **(B)** Temporal expression windows of the promoters used to manipulate MUS and OPL2 during the GMC-to-GC transition. C, F, I) Schematic of MUS (C), OPL2 (F), and TM-OPL2 (I). (D, E, G, H, J, K) Representative images of MUS-VENUS (green; D, E), OPL2-VENUS (magenta; G, H), or TM-OPL2-VENUS (magenta; J, K) expressed under the indicated pre-GMC division promoters (MUTEpro, D and J; TMMpro, G) or the FAMA promoter (E, H, K). Dotted arrows indicate the orientation of the intensity profiles. Right, mean fluorescence intensity profiles across the ventral and outer plasma membrane (n = 10 GCs) with the predominant localization pattern indicated schematically. Scale bars, 5m, all images at same scale. *Associated supplemental figure: S6*

The cytosolic IDR/scaffold protein OPL2 localized robustly to the ventral wall membrane region regardless of the promoter driving it (Fig. 2F), suggesting the protein does have intrinsic polar targeting. To subject OPL2 to cell-cycle dependent secretory flow, we fused it to the extracellular and transmembrane domains of MUS, creating TM-OPL2 (Fig. 2G). TM-OPL2 expressed before division could be maintained at the poles, whereas expression during division resembled WT OPL2 in its localization to the ventral wall membranes in maturing GCs (Fig. 2F-G, fig. S6). Membrane availability and the developmental timing of protein delivery can therefore encode spatial information and provide tools for repositioning both MUS and OPL2.

If MUS and OPL2 are not merely markers of pore position but actively define it, then altering their localization should redirect or abolish pore formation. Directing OPL2 to cell junctions produced heart-shaped stomata where GCs largely retained their kidney shape and formed pores, though at eccentric edge positions (Fig3E-F; fig. S7). Directing MUS to the ventral wall (4 independent FAMApro:MUS lines, fig. S7) produced a striking continuum of aberrant stomatal architectures: from morphologically normal (∼40%), to a complete inversion of sister GC orientations such that stomata resembled butterflies, with pores positioned outward (∼12%) (Fig. 3G) to uniform expansion and loss of GC polarity (18%, Fig. 3H). In the remaining ∼30% of stomata, the GCs exhibited mixed morphologies and orientations, consistent with each GC acting independently (fig. S7).

**Figure 3:**
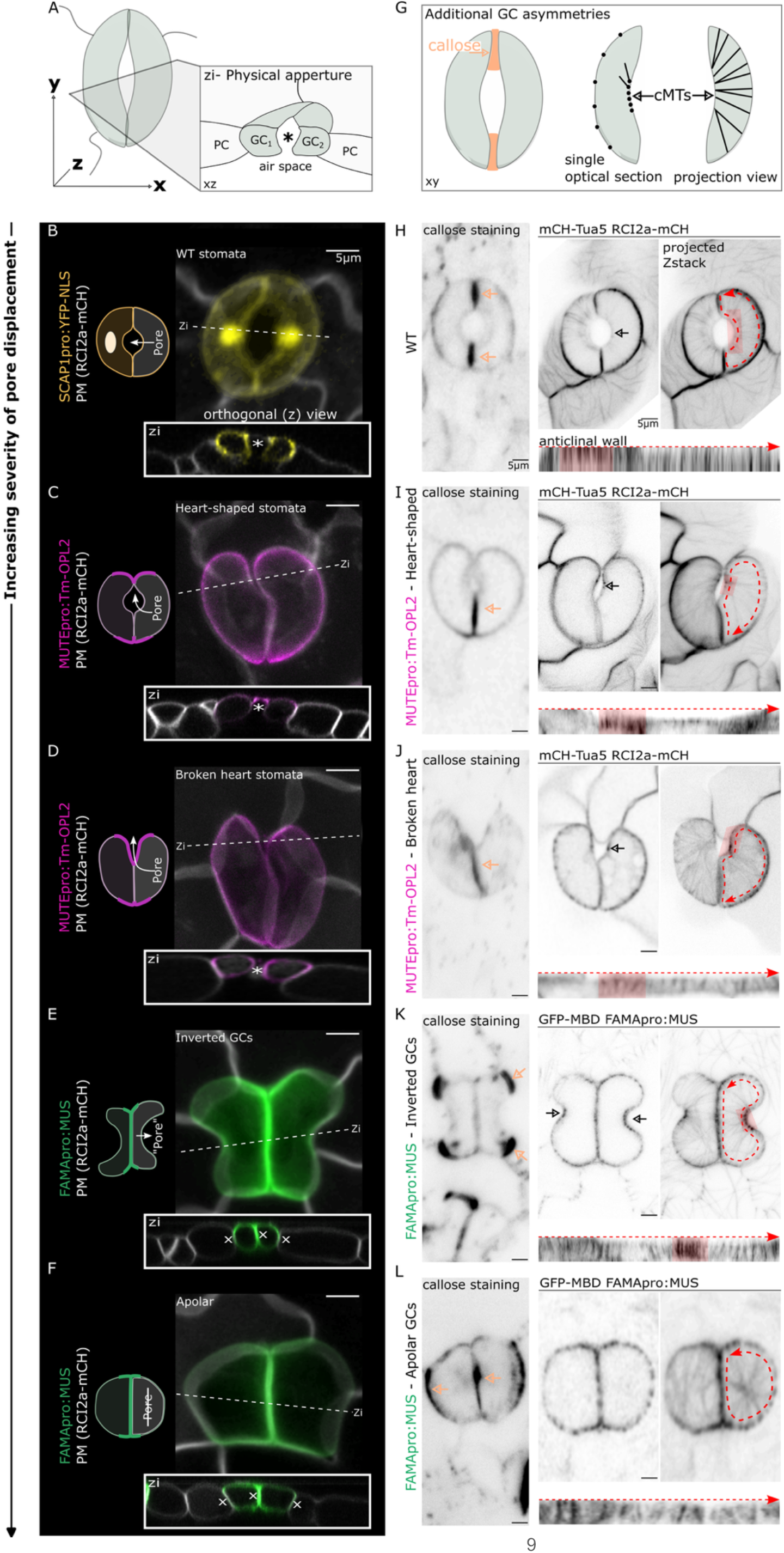
Relocalizing MUS or OPL2 results in miscoordination, reversal or elimination of GC polarity and pore positions. **(A)** Schematic of a stomatal complex in tissue context, indicating orthogonal imaging planes (Zi) and the physical aperture (*). **(B-F)** Mature stomatal complexes of WT and manipulated OPL2 and MUS lines, each with a schematic, a projected Z-stack (longitudinal), and an orthogonal (XZ) view; asterisk * indicates a visible pore. The phenotypes are presented from top to bottom in order of increasing severity of pore displacement. **(B)** WT stoma expressing YFP-NLS under SCAP1 promotor **(C to F)** Representative classes of aberrant stomatal phenotypes resulting from TM-OPL2 expression [**(C)** and **(D)**, shifted pore] or FAMApro:MUS induction [**(E)**, inverted GCs; **(F)**, apolar GCs]. ×, indicates no visible pore in XZ view. **(G)** Schematic representation of GC callose and cortical MT asymmetries. **(H to L)** Asymmetric GC marker distribution in wild type **(H)**, shifted pore [**(I)** and **(J)**], inverted GCs **(K)**, and apolar **(L)** GCs. Left, callose localization. Right, cortical microtubule and plasma membrane organization in medial (left), Z-projected (right), and resliced (bottom) views; black arrows, focus of microtubule organization; orange arrows, callose enrichment; red dotted arrow indicates orientation of the anticlinal wall relative to the maximal projection. Red shading in anticlinal wall view indicates MT foci. Scale bars, 5m, all images at same scale. *Associated supplemental figures: S7-8*

The morphological changes were striking, but it was unexpected that a single genetic manipulation could completely invert the ancient, conserved stomatal form. We therefore examined other features critical to GC form and function in the ‘rewired’ lines. In unperturbed GCs, kidney-shaped morphology is accompanied by polarized localization of cell wall and cytoskeletal components (6, 8, 20-22), and the pore is a site of complete cell separation (23). In heart-shaped and inverted stomata, callose, pectin, and microtubules shifted in absolute localization but maintained consistent orientations relative to each GC’s presumptive pore (Fig. 3I–H; figs. S7 and S8). When challenged with abscisic acid (ABA) or light, signals that change stomatal aperture, both WT and inverted GCs can change shape, suggesting that inverted GCs have some ability to perceive stimuli and mount a turgor-driven response (fig. S8). Physical aperture formation at this site, however, occurred only when a GC faced another GC, as in heart-shaped stomata or those where both GCs faced the same direction (Fig. 3; figs. S7 and S8). Taken together, the suite of phenotypes observed upon rewiring suggests that: (1) GCs can maintain individual polarity while losing sister-cell coordination; (2) polarized subcellular organization of GC walls and MTs is cell-autonomous and/or downstream of MUS and OPL2 activity; and (3) contrary to expectations from current models (6, 7), stiffened polar regions pushing against one another are not required to generate kidney-shaped GCs.

Because MUS and OPL2 appeared to regulate multiple aspects of GC form, we tested their epistatic relationships to begin building a regulatory circuit. Forced relocalization of OPL2 to the outer wall-adjacent membrane (‘outer membrane’) by MUTEpro:TM–OPL2 did not alter MUS enrichment at the cell poles (fig. S9A,B). Redirecting MUS to the ventral-wall adjacent membrane (‘inner membrane’) by FAMApro:MUS, however, profoundly affected OPL2 localization. Time-lapse imaging of FAMApro:MUS stomata revealed three distinct OPL2 responses, each correlated with a specific final stomatal morphology. If OPL2 remained at the ventral membrane, resultant GCs were oriented as in wild type (Fig. 4B). If OPL2 relocated to the outer membrane, stomata were inverted (Fig. 4C). If OPL2 was removed from the ventral membrane but failed to coalesce at the outer membrane, GCs were apolar (Fig. 4D) (Fig. 4B–D). Live cell imaging across GMC divisions enabled us to see progressive MUS accumulation at the ventral membrane accompanied by OPL2 exclusion (fig. S10A). Pseudo-timecourse analysis ordered by increasing MUS ventral enrichment revealed a progressive switch of OPL2 enrichment from the inner to the outer membrane (Fig. 4E), where OPL2 then re-established a polarized domain of comparable size. Consistently, we found an exclusion zone of >2.5 µm between MUS and OPL2 proteins, both in the wild-type ventral membrane and in the outer membrane of inverted stomata (Fig. 4F; fig. S10B,C). This relationship suggests that MUS enforces the pole-avoidance mechanism (Fig. 4F) and predicts that loss of *MUS* would lead to expansion of OPL2 along the inner membrane. OPL2 can indeed reach the poles in *mus*; however, as seen in Fig. 4G and fig. S11, OPL2 typically extends to a single pole. Wherever OPL2 accumulates – in its normal centered ventral domain, off-center in *mus*, or ectopically in the outer membrane – it forms a domain of the same length, with an intensity gradient peaking at its center (Fig. 4B, C, G; figs. S9, S11). OPL2, therefore, has some capacity to coalesce into a domain of stereotyped dimensions independently of *MUS*.

**Figure 4:**
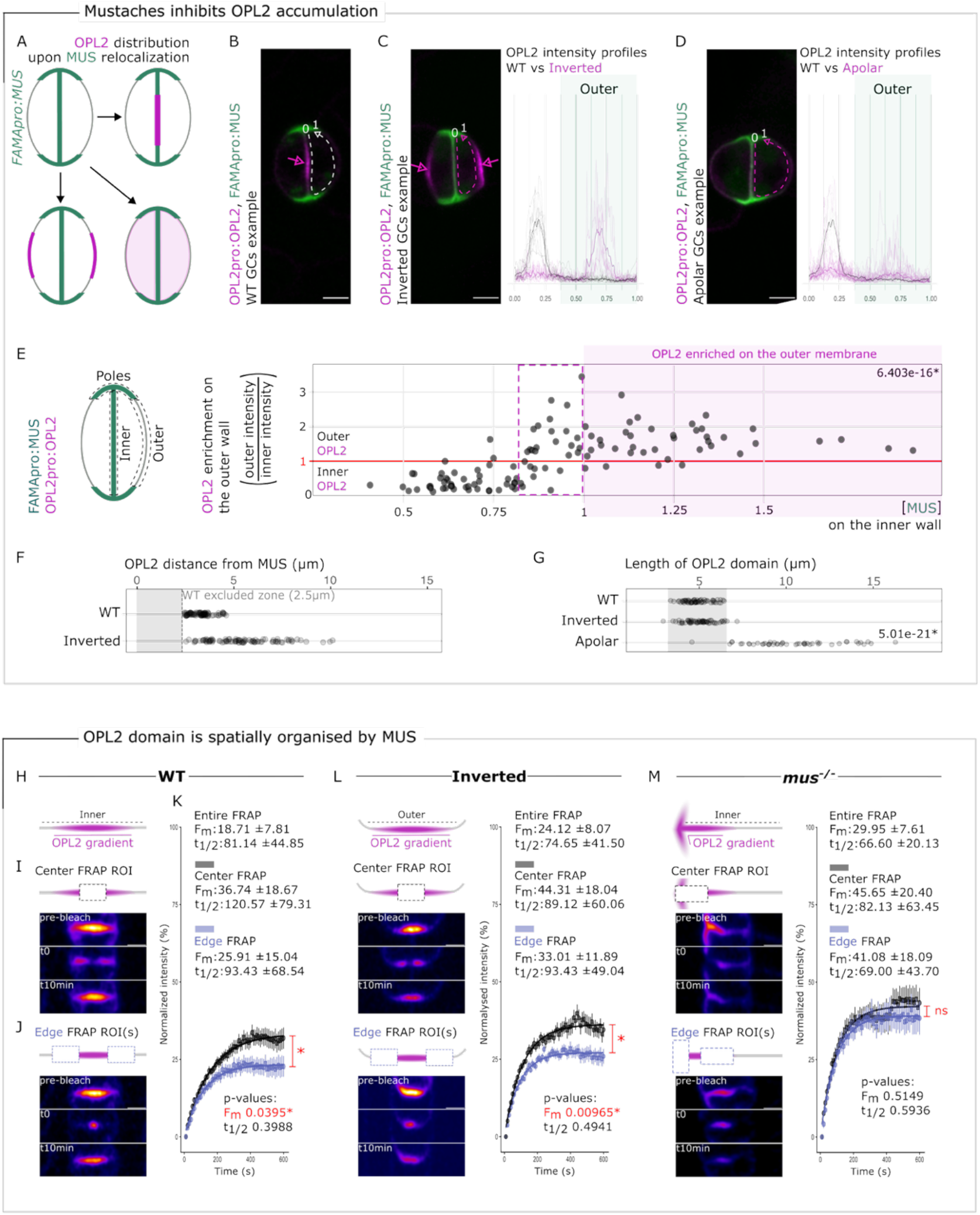
MUS directs OPL2 position and dynamics at the membrane. **(A)** Cartoon showing classes of OPL2 responses to MUS relocalization **(B to D)** Representative medial sections of stomata expressing both OPL2pro:OPL2-VENUS and FAMApro:MUS (hereafter OPL2 and MUS) for each phenotypic class: WT-like [(B), 479/1140], inverted GCs [(C), 130/1140], and apolar [(D), 203/1140]; each panel shows the image and corresponding OPL2 intensity profile (magenta), overlaid in (C) and (D) with the WT-like profile (gray, black average). **(E)** OPL2 outer-wall enrichment (outer/ inner intensity, n = 114) as a function of MUS inner-wall enrichment (inner/poles intensity). Schematic (left) defines quantified domains; purple dotted box marks the threshold above which OPL2 relocates to the outer wall. * Represents significance, Mann-Whitney - Wilcoxon rank sum test with continuity correction (MUS intensity when OPL2 intensity ratio <1 vs >1). **(F)** Distance between MUS and OPL2 reporter distributions in GCs of 10 independent 7-dpg cotyledons (WT, n = 92; inverted, n = 96). **(G)** OPL2 domain size in 10 independent cotyledons for WT (n = 64 GCs), inverted (n = 62 GCs) and apolar (n = 49 GCs). * Represents significance, Kruskal-Wallis - Dunn (Bonferroni) Post hoc test. **(H to K)** FRAP analysis of OPL2 behavior in GCs. (H) Diagram of OPL2 gradient before photobleaching. (I) and (J) Representative time series (pre-bleach, t = 0, t = 10 min) for center and edge ROIs. (K) Recovery curves (normalized fluorescence vs. time, 10-s intervals) for center [black, n = 24] and edge [blue, n = 21] ROIs, with mean ± SD, Fm, t½, and statistical comparison (* Represents significance, Mann-Whitney - Wilcoxon rank sum test with continuity correction). **(L)** FRAP analysis of OPL2pro in Inverted GCs, presented as in (H) to (K) ; center ROI, black (n = 28); edge ROI, blue (n = 26). **(M)** FRAP analysis of OPL2pro in *mus* GCs, presented as in (H) to (K); center ROI, black (n = 16); edge ROI, blue (n = 15). Scale bars, 5m, all images at same scale. *Associated supplemental figures: S9-12*

To gain insights into how OPL2, an unstructured protein lacking a transmembrane domain, can generate a defined gradient at membranes, and how OPL2 is influenced by MUS, we used FRAP to assess OPL2 dynamics in wild type, inverted, and *mus* GCs, either bleaching the entire OPL2 domain or regions of interest (ROI) at its center or edges (Fig. 4H–O; fig. S12A-C, M, N). Across all genotypes and regions, OPL2 exhibits a low Fm and slow t½ – signatures of a substantial fraction trapped in a long-lived, essentially non-exchanging state at the membrane. FRAPing the center of the wild-type OPL2 domain led to more rapid and complete recovery than FRAPing the edges (Fm: 36.7 ±18.7% vs. 25.9 ±15.0%; p = 0.040) (Fig. 4K) The same center–edge asymmetry was preserved when OPL2 was ectopically located on the outer membrane of inverted stomata (Fig. 4M; fig. S12D-F, M, N), demonstrating that these dynamics are intrinsic to OPL2 and largely independent of the membrane it occupies. In *mus* mutants, however, center–edge asymmetry was abolished (p = 0.51) and the overall mobile fraction increased (Fig. 4N, fig. S12G-I, M, N) consistent with MUS creating inhibitory domains from the poles that suppress OPL2 membrane dynamics.

A MUS-OPL2 polarity circuit may set pore location but still requires downstream effectors to translate that information into morphology. While characterizing OPL2 and MT reporter behavior, we noticed that the peak of OPL2 abundance was perfectly aligned with the focus of MTs emanating from the presumptive pore in wild-type GCs (5) (Fig. 5A; fig. S13A, and Movie S1). Genetic perturbations support a causal role for the OPL2 gradient in organizing this MT array. Flattening the gradient by increasing OPL2 levels during the GMC division (FAMApro:OPL2) abolished or delayed MT focusing, producing sparse arrays adjacent to the ventral wall (Fig. 5B). In inverted GCs where OPL2 is relocalized to the outer wall, MTs emanate from the ectopic OPL2 foci (Fig. 5C fig. S13B). In apolar GCs that fail to establish any OPL2 domains, MTs did not reorganize from their post-division transverse orientation (Fig. 5D; fig. S13B).

**Figure 5:**
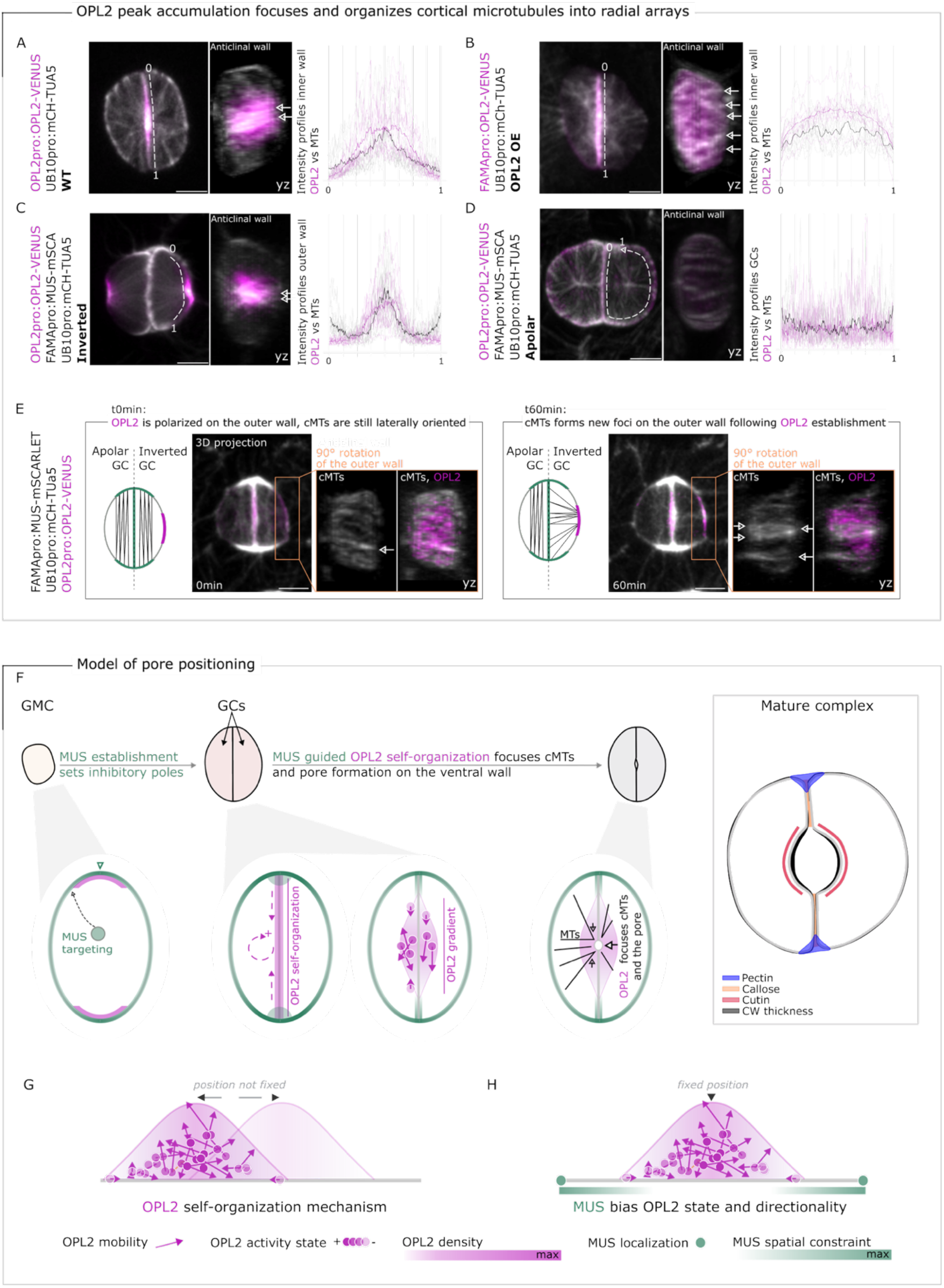
Stomatal morphogenesis involves a polarity circuit of MUS leading to OPL2 positioning and an OPL2 gradient focusing microtubules at its peak. **(A to D)** Max projections (xy and yz) and intensity profiles (n = 10 GCs) of OPL2 (magenta) and MTs (white) in WT (A, B), Inverted (C) and Apolar (D) GCs at 3dpg. OPL2 is expressed under its native promotor (A, C, D) or overexpressed with FAMA promotor (B). Dashed white line indicates position of yz plane in right image and intensity plots. **(E)** Early and late frames from time-lapse of FAMApro:MUS induced GC reorganization; The two GCs have different trajectories as shown by OPL2 (magenta) and cortical MT (grey) reorganization and cartoons **(F)** Model of pore positioning in normal GCs: Following GMC division, MUS is targeted to and establishes inhibitory poles at the ventral–outer wall junctions. Within the resulting MUS-free zone, OPL2 undergoes MUS-guided self-organization into a graded domain centered on the ventral wall. The resulting OPL2 gradient focuses cortical microtubules (MTs) at its peak, which in turn directs pore formation at this position. In the mature stomatal complex, this early polarity is reflected in the differential distribution of wall components: pectin and callose accumulate at the cell poles, while cutin and increased wall thickness mark the pore margins. **(G-H)** Most parsimonious biophysical models for behavior of OPL2 and MUS: (G) OPL2 can assemble into a gradient of stereotyped dimensions at the membrane, independent of MUS; (H) native or ectopic MUS defines the region in which OPL2 can self-assemble. Arrow length and direction reflect OPL2 movement; cytosolic exchange and repulsion by MUS (green) from cell junctions are indicated.*Associated supplemental figures: S13-14*

OPL2 domain establishment and MT reorganization normally occur within ∼1 hour of wild-type GMC division, making their temporal relationship difficult to resolve (fig. S13A). The OPL2 relocalization that occurs in inverted GCs provided a useful temporal separation of these events. Time-lapse imaging of rewired stomata in which the two daughter GCs independently adopted different outcomes showed that radial MT arrays emerged only after a stable OPL2 domain had formed, and always in register with that domain (Fig. 5E and Movie S2). Neither OPL2 domain size nor position in either wild-type or inverted GCs, however, was affected by oryzalin treatments that depolymerized MTs (fig. S13C,D). These data suggest that OPL2 localization is a cause, not a consequence of MT organization and GC shape, and that OPL2 is both necessary and sufficient to focus cortical MTs in guard cells, regardless of which membrane it occupies.

## DISCUSSION

Together, our results define core elements of pore positioning and the coupled asymmetries that generate the overall symmetry of stomata (Fig. 5F). MUS localization establishes inhibitory poles at the ventral–outer wall junction in newly formed GCs, generating conditions unfavorable for OPL2 accumulation. Outside of the MUS zones, OPL2 forms a concentration gradient peaking in the center of each ventral wall that focuses cortical MT reorganization. MTs in turn direct asymmetric cell wall deposition that drives GC shape, likely by constraining relative expansion in the pore region (Fig. 5F). We used our quantitative and dynamic OPL2 and MUS data to evaluate eight plausible general models for their distinct localization in GCs (see methods and fig. S14). An integrated model consistent with all current data is that OPL2 interconverts between an individual state characterized by slow or directionally biased movement and a fast-diffusing/self-associating state (*Df*); this latter state self-organizes into a centered, mass-conserving domain (Fig. 5G). MUS inhibits *Df*, resulting in removal of OPL2 from polar regions in WT, and from the inner wall in FAMApro:MUS-driven “rewired” stomata (Fig. 5H).

MUS and OPL2 are plant-specific molecules, but the pore-positioning strategy they implement parallels polarity mechanisms in other organisms. The behavior of OPL2 at membranes within MUS-defined boundaries is predicted by mass-conserving reaction-diffusion (McRD) systems, where an upstream template confines a downstream component, and two symmetric exclusion boundaries spontaneously produce a gradient peaking at domain center (24, 25). The pole-avoidance logic used to build stomata parallels centering strategies used in microbes – including the Min oscillator that positions the division site in *E. coli* and the Pom1-Mid1 pathway that centers cytokinesis in fission yeast (26-28). Stomata, however, have an additional requirement: two sister cells must coordinate such that their shared wall hosts a single centered pore. The elegant solution is to link the spatial distribution of MUS to the temporal event of GMC cytokinesis. This temporal-to-spatial conversion is conceptually related to cell-cycle-gated PAR recruitment in *C. elegans* (29), but here timing is coupled to plant-specific modes of secretion rather than cell-cycle dependent kinase signaling.

The MUS–OPL2–MT circuit we have defined here opens numerous questions and opportunities. MUS and OPL2 are co-expressed earlier in the stomatal lineage, yet their mutual exclusion is specific to guard cells (15, 16), implying that GC-specific factors license the antagonistic circuit, a reasonable idea given the massive transcriptome and proteome changes associated with acquiring GC identity (10, 30). The molecular basis of MUS-mediated OPL2 exclusion (as posited by our models), is currently unresolved; MUS is a leucine-rich repeat receptor-like kinase, but it lacks residues considered essential for canonical kinase activity, leaving open the mechanism by which it creates inhibitor zones. The ability to rewire the circuit — redirecting MUS or OPL2 to ectopic membranes and producing a predictable phenotypic spectrum from shifted pores to inverted or pore-less stomata — provides a genetically tractable platform for dissecting how polarity information is transduced into cell wall architecture and how sister cells achieve morphological coordination through shared developmental history. The inverted butterfly and apolar configurations allow single-cell polarity, bilateral coordination, and pore formation to be evaluated independently, offering a powerful entry point into general principles of pattern formation and the ability to test specific hypotheses (31) about the biophysical requirements to make functional stomata.

## MATERIALS AND METHODS

### Plant materials and growth conditions

Arabidopsis thaliana accession Columbia-0 (Col-0) was used as the wild-type reference. All transgenic lines are listed in **table S1**. Seeds were sterilized for 3–6 h using chlorine gas and germinated on half-strength Murashige and Skoog (½ MS) medium containing 0.7% (w/v) plant agar and 0.5% (w/v) sucrose (pH 5.7). Seedlings were grown under long-day conditions at 22°C. Cotyledons were imaged between 3 and 14 days post-germination (dpg) while maintained on ½ MS medium. For time-lapse imaging, young true leaves were collected from soil-grown plants at 21–42 dpg grown under long-day conditions.

Inducible overexpression of MUTE (iMUTE, plasmid described in (32) was achieved by growing seedlings for 3 dpg on ½ MS medium supplemented with 10 µM estradiol. For Oryzalin treatment, 3-dpg seedlings were incubated in liquid ½ MS + 0.75% sucrose supplemented with 20 µM Oryzalin or an equivalent volume of DMSO for 2 h. Seedlings were kept in their mounting solution when placed on slides. Other plant species used in this study (listed in **table S1**) were grown directly in soil, and cotyledons or true leaves were imaged depending on developmental stage.

### Cloning and Agrobacterium-mediated transformation

Constructs for plant transformation were generated using the GreenGate system with the pGreenII backbone (33). Module C constructs containing MUSTACHES cDNA or MUSTACHES–OPL2 chimeras were assembled in two steps. First, the MUSTACHESext–tm fragments were amplified using Phusion polymerase and GreenGate-specific primers (**table S2**) and cloned into pUC19 entry modules via BsaI restriction. The MUSTACHESint region or OPL2 was then inserted by Gibson assembly. Final constructs (**table S3**) were produced by assembling the corresponding entry modules following the GreenGate procedure (33). All plasmids were verified by whole-plasmid sequencing and introduced into *Agrobacterium tumefaciens* strain GV3101 by electroporation. Wild-type Col-0 homozygous *mus* plants, or other transgenic lines were transformed using the floral dip method (34). Transformed seedlings were selected on ½ MS plates with 0.8 % plant agar but without sucrose and the respective antibiotic/herbicide (7.5 mg/L sulfadiazine, 50 mg/L kanamycin, 50 mg/L hygromycin, 15 mg/L phosphinothricin) as listed in Table S1.

### Imaging and tissue staining

Images were acquired on a Leica Stellaris confocal microscope (LAS X software, leica-microsystems.com) equipped with a white light laser and HyD detectors, using a 40× NA 1.1 water objective. Excitation and emission settings were chosen to match fluorophore specifications as closely as possible while minimizing bleedthrough and autofluorescence. For microtubule time-lapse imaging, true leaves were mounted in ½ liquid MS (0.5% sucrose) between two coverslips separated by a 0.5-mm silicone spacer that provided mechanical support. Images were acquired every 20 min (when combined with OPL2 imaging) to 30 min, over a period of 4 to 8 h. OPL2 time-lapses were performed on both cotyledons and true leaves using the same mounting strategy, with 10-min time intervals over a 4-h duration.

FRAP experiments were performed on first true leaves mounted in ½ liquid MS (0.5% sucrose) between two coverslips, using a Leica SP8 inverted confocal microscope equipped with a 514-nm argon laser (65 mW) for imaging and bleaching, and controlled by Leica LAS X software (Stanford Cell Sciences Imaging Facility). Images were acquired using a 40× HC PL APO CS2 objective lens (NA 1.30; working distance 240 μm) with the confocal pinhole set to 5 Airy units (AU) to compensate for sample drift. Photobleaching was performed using 30 iterations at 100% laser transmission. FRAP was recorded continuously using the same parameters as the pre-bleach acquisition: 2% laser power for 600 s, with a 10-s delay between frames. Fluorescence intensities were measured over time within the bleached ROI and corrected for background fluorescence and acquisition photobleaching using unbleached reference regions (the ventral wall of unbleached neighboring stomata). Normalized recovery curves were generated relative to pre-bleach fluorescence intensity. Four to ten ROI were acquired per leaf in two separate acquisitions, with at least eight leaves used per condition across three independent experiments.

Callose deposition was visualized in living stomatal complexes using aniline blue staining (adapted from (35). Seedlings were incubated in 0.05% (w/v) aniline blue in 0.07 M K2HPO4 (pH 8.5) for 30 min to 1 h. For true leaves of non-Arabidopsis species, the incubation time was extended to 90 min.

Immunolocalization of pectins was performed on hand-cut paradermal leaf sections (adapted from (35). Tissues were fixed in 8% (w/v) paraformaldehyde in PEM buffer (50 mM PIPES, 5 mM EGTA, 5 mM MgSO4, pH 6.8) for 45 min, washed, and subjected to enzymatic digestion with 1% (w/v) cellulase for 90 min (supplemented with 2% macerozyme for 40 min in wild-type samples). After permeabilization in PBS containing 0.5% Triton X-100 and 2% DMSO, sections were blocked in PBS with 2% (w/v) BSA, incubated overnight with JIM5 antibody (1:40), washed, and labeled with alexa488-conjugated goat anti-rat IgG antibody (1:40) 3hours at 37°C. Samples were mounted in glycerol/PBS (2:1, v/v) containing 0.5% p-phenylenediamine.

COS488 labeling was performed as described in (36). Seedlings were incubated in COS488 (1:1000) diluted in 25 mM MES (pH 5.7) for 20 min, rinsed in MES, and mounted in MES:glycerol (1:1).

Cutin autofluorescence was excited by a 405-nm fixed diode laser, and emission was collected in the 450-500 nm range in live tissues. Z-stacks were acquired to capture the full depth of the guard cell complex.

### Stomatal opening and closure assay

Cotyledons from 6 dpg seedlings were first incubated in light for 2.5 h in a solution containing 20 mM KCl, 1 mM CaCl2, and 5 mM MES-KOH (pH 6.15) to allow stomatal opening. Stomatal closure was then induced by incubating half of the seedlings in the same solution supplemented with 50 µM Abscisic acid (ABA) in the light for an additional 2.5–3 h. The other half was kept without ABA under the same conditions (37).

### Image quantification

Fluorescence intensity profiles along the guard cell cortex were extracted in FIJI/ImageJ (fiji.sc, version 2.14.0/1.54f, (38) using 3-pixel-wide line scans tracing the entire cell perimeter. Line scans were oriented consistently, beginning at one pole of the guard cell and passing first through the inner membrane and then the outer membrane. Fluorescence values were normalized to the cytosolic background, measured from a region corresponding to 40–60% of the cytoplasmic area. For each cell, the intensity at every position along the line was divided by the background value, and normalized profiles from ten cells were aligned on a common axis. At least 3 independent cotyledons were used.

Spanning wall dimensions/callose domains were measured manually from a central optical section of each Z-stack, corresponding to the plane with the largest stomatal perimeter, using a straight line in FIJI drawn parallel to the inner wall. Spanning membrane length and stomatal length (measured as the distance between the two poles of the guard cell pair) were quantified for each stoma, and pore length was calculated as: stomatal length minus combined spanning wall length. The same strategy was used to measure cell wall stubs in the *map65-3* mutant, with the exception that stub size corresponded either to the entire callose-stained domain (when no pore was formed) or to the maximum distance between the cell pole and the tip of the pore (when a pore was formed).

For guard cell shape analysis after opening or induced closure, cell contours were obtained from manually defined ROIs where we measured the left guard cell from each wild-type stomatal complex (right GC for inverted ones) to be able to superimpose contours (as in **fig. S8**). Maximum-intensity projections of Z-stacks were binarized, and objects were segmented using the FIJI “Analyze Particles” plugin. Data processing and plotting were performed in RStudio (r-project.org). For each segmented cell, three parameters were extracted: maximum guard cell length, guard cell width at the cell midpoint (“minimum GC width”), and guard cell width at one-third of the maximum length (“top lobe width”).

Intensity ratios in **fig. S6** were calculated from mean fluorescence values measured along manually drawn lines tracing either the outer or inner wall/membrane (pole to pole) of individual guard cells. **In Fig. 4E**, ‘pole’ intensity corresponds to the average signal from both MUS pole domains. For OPL2 measurements, the outer wall was defined by the MUS signal (from the edge of the MUS ‘north’ pole domain to the edge of the MUS ‘south’ pole domain), whereas the inner wall was defined by the cell poles.

Polar domain size has been measured in various ways in the literature, but no consensus has been reached on an optimal approach. In this study, we defined a polar domain as a continuous region of detectable fluorescence with “ends” exhibiting at least one-third maximal intensity. Intensity profiles were extracted from a 3-pixel-wide line drawn across each polar domain. The maximum intensity value of the domain was divided by three, and domain size was determined by measuring the distance between the points where the intensity profile crossed this threshold (maximum intensity/3). For consistency, the same OPL2pro line was used for the different engineered polarity manipulations. OPL2pro:OPL2-VENUS was introduced into *mus* mutant background separately. The same approach was used to quantify distances between polar domains.

Stomata were counted in seedlings at 7 days post-germination, with ten seedlings analyzed per line and two independent lines per construct. For each seedling, one abaxial region of 465.52 µm^2^ (containing 17 to 49 stomata) was randomly selected. Stomata were then classified by morphology (wild type, inverted GCs, apolar, heart-like, and mixed forms, as shown in **fig. S7**).

### FRAP configuration and analysis (Fig. 4H-O, fig. S12)

Three complementary FRAP configurations were used to characterize OPL2 dynamics: (1) whole-domain FRAP, in which the entire OPL2 cortical crescent was bleached to measure fluorescence recovery resulting from exchange with the external cytoplasmic or membrane-associated pool; (2) internal FRAP, in which a small ROI positioned within the OPL2 domain was bleached to assess internal molecular rearrangement and intradomain mobility; and (3) edge FRAP, in which two identical ROIs positioned at the boundary of the OPL2 domain were bleached to evaluate lateral diffusion and directional recovery from surrounding regions. For each configuration, we performed the following:

- *Kymograph*
- *Fitting curve:* I(t) = A(1 − e−kt)

*where I(t) is the fluorescence recovery at time t, A is the maximum recovery (mobile fraction), k is the recovery rate constant, and t is time*.

- *Mobile fraction (F*m):

Fm = [(Ie − I0) / (Ii − I0)] × 100%

where Ie is the mean intensity at the endpoint (t = 600 s), Ii is the mean initial intensity before bleaching, and I0 is the mean intensity at t0.

- *Recovery half-time (t*1/2):

Determined from fitting curves applied to normalized recovery curves, as the time required to reach 50% of the maximal fluorescence recovery.

- *Recovery rate constant (k):* t1/2 = ln2 / k
- *Mobile fraction index (F*m *index) / diffusion index (D index):*

Estimates of cytosolic exchange versus lateral diffusion, for both center and edge recovery.

Fm index = Fm (edge or center) / Fm (whole domain)

D index = t1/2 (edge or center) / t1/2 (whole domain)

### Comparative evaluation of candidate MUS–OPL2 mechanistic models

To ask how MUS and OPL2 generate two spatially segregated, polarized membrane domains, we scored eight candidate mechanistic models (**fig. S14**) representing classical polarity/morphogen mechanisms, two additional physical models, and a falsifiable MUS-independent null against 14 discrete predictions drawn from our imaging, genetic, and FRAP data (**fig. S14**). These predictions covered domain geometry and dosage, MUS gain- and loss-of-function, the spatial gap between domains, protein trafficking, and centre-versus-edge FRAP kinetics. Each model was scored per criterion as compatible (“Fit”, =1), compatible only with an added untested assumption (“Partial”, =0.5), or contradicted (“No fit”, =0), and tallied (max 14) as a heuristic ranking, not a statistical test (**fig. S14**).

| # | Model | Core proposal | Fit score |
|---|---|---|---|
| A | Picket-fence corral | MUS locally raises cortical/lipid packing density at the boundary, increasing the trapping rate there. | 10.5/14 |
| B | Capture-and-clearance | MUS converts mobile OPL2 to an immobile, marked state at the edge, which is then cleared. | 10/14 |
| C | Decoupled source + edge trap | Polarized delivery to the domain center, mechanistically independent of a separate edge-immobilization step. | 11/14 |
| D | Advective transport | Directed cortical flow sweeps OPL2 toward the center; untransported material strands at the periphery. | 7/14 |
| E | Bistable front (biased by MUS) | OPL2 interconverts between a low- or directed-diffusing state and a fast-diffusing, self-associated state; MUS locally biases this transition | 13/14 |
| F | Direct binding / tether | MUS directly and non-catalytically binds OPL2 at the boundary, forming a stable complex. | 8.5/14 |
| G | Steric exclusion | MUS receptor clusters crowd the pole membrane, excluding OPL2 by volume competition alone. | 3/14 |
| H | MUS-independent null | OPL2 self-limits by intrinsic nucleation and turnover alone; MUS plays no role. | 6/14 |

The 14 criteria (**fig. S14**) spanned five categories:

- Domain geometry and dosage (stereotyped domain length, a peak that is always central, a saturating response at high overexpression, and domain formation by OPL2 alone in heterologous *N. benthamiana* cells).
- MUS gain/loss-of-function (dose-dependent clearance of an established domain, occasional repolarization failure, one-directional coupling, and loss of control when a membrane-locked OPL2– MUS-TM fusion occupies the MUS pole).
- Spatial relationship (an obligate, size-scalable ≥2.5 µm gap between the two domains).
- Trafficking (absence of puncta, and a diffuse cytoplasmic rather than immobilized fate for displaced OPL2).
- FRAP kinetics (genotype-dependent whole-domain Fm, a centre>edge Fm asymmetry in WT, inverted, and enlarged-OPL2 stomata, no statistically significant centre-versus-edge difference in t½ in any genotype, and loss of the Fm asymmetry specifically in *mus*).

Most models were clearly refuted, each on distinct grounds: B predicts an immobilized, punctate clearance product that is not observed; D requires that under each condition (WT, *mus*-/- and inverted) that there is faster FRAP recovery at the center than at the edge, and this is not seen; and H is contradicted by every MUS gain- and loss-of-function result, including the OPL2–MUS-TM titration. G has no mechanism for a graded peak or dose-response. A and F reproduce the center-high/edge-low mobile-fraction (Fm) pattern but not the obligate ≥2.5 µm MUS–OPL2 gap, which a contact-range fence or bound complex should not leave. Model C (central delivery source plus an independent edge trap) is hard to reconcile with the consistent shape of the OPL2 gradient whether or not MUS is present: a genuinely MUS-independent delivery source would still need to be directed to the inner-wall center by some other mechanism, and it is unlikely that two unrelated polarization pathways would converge on an identical outcome.

E (OPL2 interconverting between a low- or directed-diffusing state and a fast-diffusing, self-associated state, with MUS locally biasing this transition) is not contradicted by any existing data and explains domain formation in the complete absence of MUS, the incomplete and dose-insensitive relocalization-failure rate, and the cytoplasmic, non-punctate fate of cleared OPL2. However, it works only if the state switch itself carries a directional bias that moves OPL2 toward the domain center, and not merely a change in mobility.

Thus, there is not a perfect model, but variations of E provide the basis for a working model. We envision this in layers. In the first layer (intrinsic, MUS-independent): OPL2 self-associates cooperatively wherever it accumulates, setting a stereotyped, self-centering domain length independent of genotype — explaining WT, *mus*, inverted, TM-OPL2, and the saturating response to extreme OPL2 overexpression. Layer 2 (graded, one-directional MUS field): MUS imposes a field that decays with distance from the poles and dose-dependently shifts local OPL2 mobility or direction, with no return coupling from OPL2 to MUS. This reproduces the scalable MUS–OPL2 gap and the center>edge Fm asymmetry, which is lost in *mus*. Layer 3 (local bistable switch, gated by Layer 2): the local mobile⇌immobile or undirected⇌directed transition behaves as a bistable ON/OFF switch, not a continuous one. Bistable and graded descriptions apply at different scales, not competing ones. Locally, each patch of membrane behaves as a two-state switch; but because the Layer-2 MUS field itself decays continuously with distance from the pole, different positions along the domain cross that local threshold at different distances from MUS. Summed across the whole domain, this produces the smoothly graded, center-to-edge spatial profile we measure.

*Confidence assessment in elements of the MUS–OPL2 circuit model:* High confidence: *(i)* OPL2’s stereotyped, self-centering domain-forming capacity is intrinsic and MUS-independent. *(ii)* The MUS–OPL2 coupling is one-directional. *(iii)* The center/edge Fm asymmetry specifically requires MUS. Medium confidence: MUS most likely acts through a graded, distance- and dose-dependent influence on local OPL2 behavior rather than through a static, contact-range physical barrier or a catalytic clearance reaction.

Working hypothesis: the local transition is bistable rather than continuously graded and involves either a change of OPL2 mobility or directionality or both. What remains open is the molecular identity of the OPL2 state that switches (a change in oligomeric/assembly state, in membrane affinity, or both), whether it involves a loss of mobility or a directional reorientation of OPL2 movement, and the biochemical nature of the MUS-derived field itself (a diffusible signal versus a locally modified property of the membrane or cortex).

### Statistical analysis

Statistical analyses were performed in R (version 4.5.1) using the RStudio interface (version 2025.09.2+418) and the packages ggplot2, lme4, car, multcomp, dplyr, tidyverse, tidyr, dunn.test, FSA, minpack.lm, and means. Graphs were generated in R/RStudio and customized in Inkscape (https://inkscape.org). In box plots, the middle horizontal line represents the median, and the bottom and top of each box represent the 25th and 75th percentiles, respectively. Whiskers extend to a maximum of 1.5 times the interquartile range (IQR), excluding data beyond this range; for ranges under 1.5 × IQR, whiskers represent the minimum and maximum values.

To assess whether a biologically relevant minimal spanning wall (SW) size could be detected in wild-type (WT) samples, we analyzed spanning wall measurements of single cells at 3, 7, and 14 days post-germination. Spanning wall size was defined as the average of the left and right measured lengths for each cell. We specifically tested whether a threshold of 3.5 µm could plausibly arise from the WT size distribution, using a bootstrap-based one-sample probability test. Briefly, individual WT spanning wall sizes were resampled with replacement 10,000 times, and the proportion of bootstrap samples in which the spanning wall size was ≤3.5 µm was calculated. This proportion was reported as a p-value, representing the probability that a randomly drawn WT spanning wall would fall below the threshold. Statistical significance was considered at p < 0.05, indicating that a cell size ≤3.5 µm is unlikely to occur in WT, supporting the presence of a minimal size. The observed WT distribution, bootstrap-derived probabilities, and descriptive statistics (mean, median, and minimum) are summarized in a table for clarity (**table S4**).

Spanning wall ratios between *Ath* lines were compared using Levene’s test for homogeneity of variance, centered on the mean (**table S4**). For size comparisons (**table S4**) and inner/outer intensity ratios (**table S5**), normality was not verified; these were therefore compared using a Kruskal-Wallis test with Dunn’s post hoc analysis (Bonferroni method). Guard cell size parameters were compared between conditions using a Kruskal-Wallis test with Dunn’s post hoc analysis (Bonferroni method) (**table S6**). MUS intensity ratios (inner/outer) were compared between the “OPL2-low outer/inner intensity (<1)” and “OPL2-high outer/inner intensity (>1)” groups using a Wilcoxon rank-sum test with continuity correction. OPL2 domain sizes were compared between conditions using a Kruskal-Wallis test with Dunn’s post hoc analysis (Bonferroni method). FRAP parameters were compared between the domain center and edge for each condition, and between conditions, using the Mann–Whitney/Wilcoxon rank-sum test with continuity correction (**table S7**).

## ACKNOWLEDGEMENTS

We thank members of the lab, particularly Macy Vollbrecht, Mabel Mathew and Aimee Uyehara for discussions and comments on the manuscript, Fridtjof Brauns (MPI-PKS) and participants of KITP’s 2026 morphogenesis program for theory discussions and Sid Shaw (IU) for FRAP and microtubule insights.

## AUTHOR CONTRIBUTIONS

AL conceived the ideas, contributed to experimental designs, investigation, analyses, data interpretation, and writing. ESW provided OPL2 materials. DCB supervised the project and contributed to writing and organization of the manuscript.

## DATA AND MATERIALS AVAILABILITY

All data required for the conclusions of the manuscript are available in the manuscript or supplemental materials.

## AI USE

Claude Sonnet was used to format materials.

## REFERENCES

1. J. M. Van Norman, Asymmetry and cell polarity in root development. Developmental Biology 419, 165–174 (2016).

2. V. V. Isaeva, N. V. Kasyanov, Symmetry Transformations in Metazoan Evolution and Development. Symmetry 13, 160 (2021).

3. P. Kenrick, P.R. Crane, The Origin and Early Diversification of Land Plants. Smithsonian Institution Press, ISBN 1560987294. 136, 213–220 (1999).

4. J. A. Nadeau, F. D. Sack, Stomatal development in Arabidopsis. Arabidopsis Book 1, e0066 (2002).

5. J. R. Lucas, J. A. Nadeau, F. D. Sack, Microtubule arrays and Arabidopsis stomatal development. Journal of Experimental Botany 57, 71–79 (2006).

6. Y. Rui, Y. Chen, H. Yi, T. Purzycki, V. M. Puri, C. T. Anderson, Synergistic Pectin Degradation and Guard Cell Pressurization Underlie Stomatal Pore Formation. Plant Physiol. 180, 66–77 (2019).

7. R. Carter, H. Woolfenden, A. Baillie, S. Amsbury, S. Carroll, E. Healicon, S. Sovatzoglou, S. Braybrook, J. E. Gray, J. Hobbs, R. J. Morris, A. J. Fleming, Stomatal Opening Involves Polar, Not Radial, Stiffening Of Guard Cells. Current Biology 27, 2974–2983.e2 (2017).

8. Tsakali, I.-C. Asitzoglou, V. Basdeki, V. Podia, I.-D. S. Adamakis, E. Giannoutsou, K. Haralampidis, The Role of PME2 and PME3 in Arabidopsis Stomatal Development and Morphology †. Biol. Life Sci. Forum 2022, 11, 36 (2021).

9. C.S. Gager, Fundamentals of botany. P. Blakiston’s Son & Co (1916).

10. S.-K. Han, X. Qi, K. Sugihara, J. H. Dang, T. A. Endo, K. L. Miller, E.-D. Kim, T. Miura, K. U. Torii, MUTE Directly Orchestrates Cell-State Switch and the Single Symmetric Division to Create Stomata. Developmental Cell 45, 303–315.e5 (2018).

11. Liu, A. Mair, J. L. Matos, M. Vollbrecht, S.-L. Xu, D. C. Bergmann, bHLH transcription factors cooperate with chromatin remodelers to regulate cell fate decisions during Arabidopsis stomatal development. PLoS Biol 22, e3002770 (2024).

12. M.-C. Caillaud, P. Lecomte, F. Jammes, M. Quentin, S. Pagnotta, E. Andrio, J. De Almeida Engler, N. Marfaing, P. Gounon, P. Abad, B. Favery, MAP65-3 Microtubule-Associated Protein Is Essential for Nematode-Induced Giant Cell Ontogenesis in Arabidopsis. The Plant Cell 20, 423–437 (2008).

13. C. M. McMichael, G. D. Reynolds, L. M. Koch, C. Wang, N. Jiang, J. Nadeau, F. D. Sack, M. B. Gelderman, J. Pan, S. Y. Bednarek, Mediation of Clathrin-Dependent Trafficking during Cytokinesis and Cell Expansion by Arabidopsis STOMATAL CYTOKINESIS DEFECTIVE Proteins. The Plant Cell 25, 3910–3925 (2013).

14. S. Keerthisinghe, J. A. Nadeau, J. R. Lucas, T. Nakagawa, F. D. Sack, The A rabidopsis leucine-rich repeat receptor-like kinase MUSTACHES enforces stomatal bilateral symmetry in A rabidopsis. The Plant Journal 81, 684–694 (2015).

15. E.-S. Wallner, L. Dolan, D. C. Bergmann, Arabidopsis stomatal lineage cells establish bipolarity and segregate differential signaling capacity to regulate stem cell potential. Developmental Cell 58, 1643–1656.e5 (2023).

16. E.-S. Wallner, A. Mair, D. Handler, C. McWhite, S.-L. Xu, L. Dolan, D. C. Bergmann, Spatially resolved proteomics of the Arabidopsis stomatal lineage identifies polarity complexes for cell divisions and stomatal pores. Developmental Cell 59, 1096–1109.e5 (2024).

17. C. Rodriguez-Furlan, R. Campos, J. N. Toth, J. M. Van Norman, Distinct mechanisms orchestrate the contra-polarity of IRK and KOIN, two LRR-receptor-kinases controlling root cell division. Nat Commun 13, 235 (2022).

18. N. Geldner, J. Friml, Y.-D. Stierhof, G. Jürgens, K. Palme, Auxin transport inhibitors block PIN1 cycling and vesicle trafficking. Nature 413, 425–428 (2001).

19. G. Jürgens, MEMBRANE TRAFFICKING IN PLANTS. Annu. Rev. Cell Dev. Biol. 20, 481–504 (2004).

20. S. Amsbury, L. Hunt, N. Elhaddad, A. Baillie, M. Lundgren, Y. Verhertbruggen, H. V. Scheller, J. P. Knox, A. J. Fleming, J. E. Gray, Stomatal Function Requires Pectin De-methyl-esterification of the Guard Cell Wall. Current Biology 26, 2899–2906 (2016).

21. L. Zhao, F. D. Sack, Ultrastructure of stomatal development in Arabidopsis (Brassicaceae) leaves. American J of Botany 86, 929–939 (1999).

22. W. R. Eisinger, V. Kirik, C. Lewis, D. W. Ehrhardt, W. R. Briggs, Quantitative Changes in Microtubule Distribution Correlate with Guard Cell Function in Arabidopsis. Molecular Plant 5, 716–725 (2012).

23. B. Roberts, What Are Stomates and How Do They Work? AUF 16, 331–331 (1990).

24. M. C. Wigbers, F. Brauns, C. Y. Leung, E. Frey, Flow Induced Symmetry Breaking in a Conceptual Polarity Model. Cells 9, 1524 (2020).

25. T. Burkart, M.C. Wigbers, L. Würthner, E. Frey, Control of protein-based pattern formation via guiding cues. Nat Rev Phys 4, 511–527 (2022).

26. S. G. Martin, M. Berthelot-Grosjean, Polar gradients of the DYRK-family kinase Pom1 couple cell length with the cell cycle. Nature 459, 852–856 (2009).

27. J. B. Moseley, A. Mayeux, A. Paoletti, P. Nurse, A spatial gradient coordinates cell size and mitotic entry in fission yeast. Nature 459, 857–860 (2009).

28. S. A. Rincon, P. Bhatia, C. Bicho, M. Guzman-Vendrell, V. Fraisier, W. E. Borek, F. D. L. Alves, F. Dingli, D. Loew, J. Rappsilber, K. E. Sawin, S. G. Martin, A. Paoletti, Pom1 regulates the assembly of Cdr2–Mid1 cortical nodes for robust spatial control of cytokinesis. Journal of Cell Biology 206, 61–77 (2014).

29. K. Ng, H. Sebaa, N. Hirani, A. Chizh, Z. Messi, T. Bland, K. Sugioka, N.W. Goehring, Cell cycle oscillations in a polarity network facilitate state switching by morphogenetic cues. Sci Advances. 15;12(20):eaec3379 (2026).

30. C. B. Lopez-Anido, A. Vatén, N. K. Smoot, N. Sharma, V. Guo, Y. Gong, M. X. Anleu Gil, A. K. Weimer, D. C. Bergmann, Single-cell resolution of lineage trajectories in the Arabidopsis stomatal lineage and developing leaf. Developmental cell, 56(7), 1043–1055.e4. (2021)

31. N. Y. H. Tan, J. V. Armand, J. E. Gray, A. J. Fleming, Modelling stomatal mechanics: a critical review. New Phytologist 249, 2736–2745 (2026).

32. K.A. Davies, D.C. Bergmann, Functional specialization of stomatal bHLHs through modification of DNA-binding and phosphoregulation potential, Proc. Natl. Acad. Sci. U.S.A. 111 (43) 15585–15590 (2014).

33. Lampropoulos, Z. Sutikovic, C. Wenzl, I. Maegele, J. U. Lohmann, J. Forner, GreenGate -A Novel, Versatile, and Efficient Cloning System for Plant Transgenesis. PLoS ONE 8, e83043 (2013).

34. S. J. Clough, A. F. Bent, Floral dip: a simplified method for Agrobacterium -mediated transformation of Arabidopsis thaliana . The Plant Journal 16, 735–743 (1998).

35. D. Paraskevopoulou, N. Anezakis, E. Giannoutsou, P. Sotiriou, I.-D. S. Adamakis, The Stomata of the Katanin Mutants, fra2, lue1 and bot1. Biol. Life Sci. Forum 2021, 4, 30. (2020).

36. Y. Rui, C. Xiao, H. Yi, B. Kandemir, J. Z. Wang, V. M. Puri, C. T. Anderson, POLYGALACTURONASE INVOLVED IN EXPANSION3 Functions in Seedling Development, Rosette Growth, and Stomatal Dynamics in Arabidopsis thaliana. Plant Cell 29, 2413–2432 (2017).

37. Y. Rui, C. T. Anderson, Functional Analysis of Cellulose and Xyloglucan in the Walls of Stomatal Guard Cells of Arabidopsis. Plant Physiology 170, 1398–1419 (2016).

38. J. Schindelin, I. Arganda-Carreras, V. Frise, E, Kaynig, M. Longair, T. Pietzsch, S. Preibisch, C. Rueden, S. Saalfeld, B. Schmid, J.Y. Tinevez, D.J. White, V. Hartenstein, K. Eliceiri, P. Tomancak, A. Cardona, Fiji: an open-source platform for biological-image analysis. Nature methods, 9(7), 676–682 (2019).

