## Supplemental Figures and Tables for "A developmentally scheduled inhibitory polarity circuit centers the stomatal pore"

**This PDF contains**

Figs. S1 to S14

**Additional supplemental materials**

**Movie S1– Timelapse of cMTs and OPL2.**
Time–course analysis of cMTs and OPL2 rearrangement from GMC to pore formation. (10 min intervals)

**Movie S2– 3D projection of Inverted GCs.**
3D rotation of Inverted GCs expressing MUS–mSCARLET, mCHERRY–TUa5 and OPL2–VENUS.

**Movie S3– Timelapse of cMTs, FAMApro:MUS and OPL2.**
Time–course analysis (20 min intervals) of cMTs rearrangement (3D projection) following OPL2 establishment on the outer membrane (right GC of both stomata).

**Supplementary tables – Resources and statistics.**
Supplementary table S1, Reagent and resources
Supplementary table S2, Primers
Supplementary table S3, Plasmids
Supplementary table S4, Details of Fig. 1 statistics
Supplementary table S5, Details of Fig. 2 statistics
Supplementary table S6, Details of Fig. 3 statistics
Supplementary table S7, Details of Fig. 4 statistics

**SUPPLEMENTARY FIGURES:**

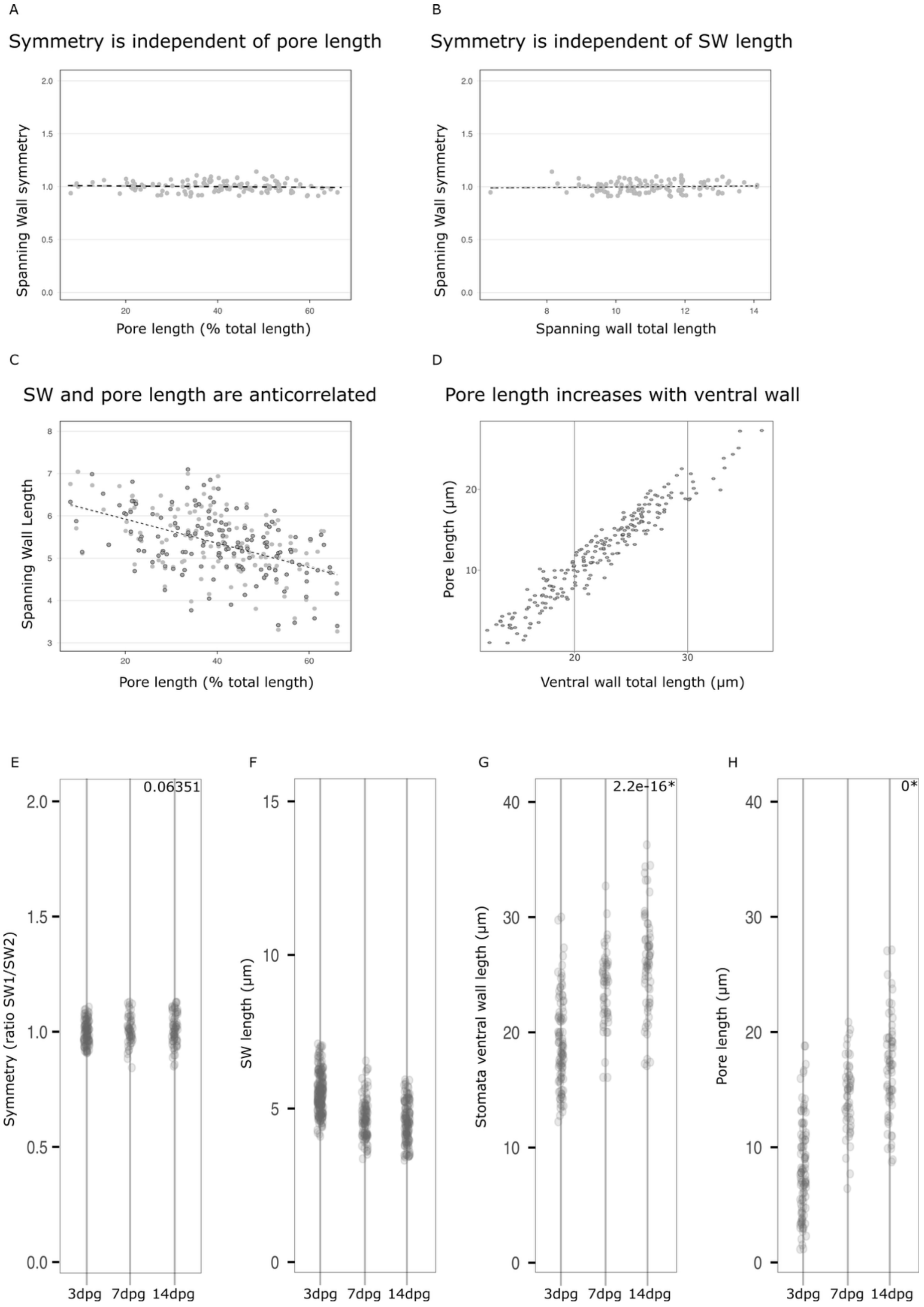

**Fig. S1 Spanning wall symmetry is invariant whereas spanning wall length decreases during stomatal maturation.**

**(A)** SW length ratio (d1/d2) as a function of pore length (% of total inner wall length, n = 100 cells, 10 cotyledons), showing constant SW symmetry across increasing pore lengths. **(B)** SW length ratio as a function of combined SW length (d1 + d2, n = 100 cells), showing symmetry is maintained independently of absolute SW length. **(C)** SW length as a function of pore length (n = 100), showing progressive SW reduction during pore expansion. **(D)** Pore length as a function of ventral wall length, showing a fast correlated pore length growth with ventral wall growth. **(E to H)** SW length ratio (E), combined SW length (F), inner wall length (G), and pore length (H) in 3, 7, and 14 dpg cotyledons (n = 100, 52 and 65 stomata, respectively). Pores are centered in young and small cells as well as in old and long ones.

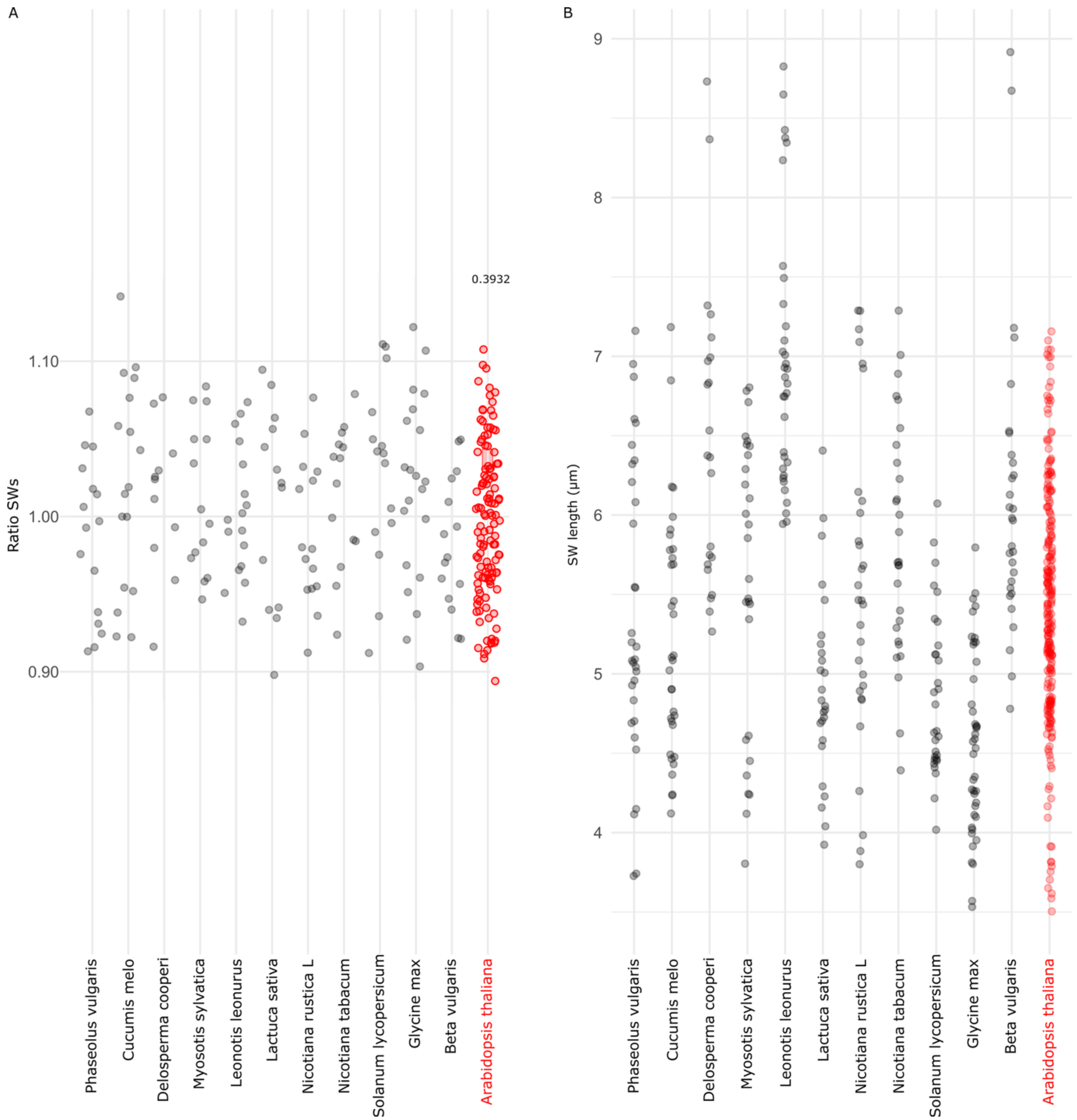


**Fig. S2 Across common angiosperm species, SW lengths exhibit minimum length and are symmetric**

**(A)** SW length ratio (d1/d2) of GCs comparing *Arabidopsis thaliana* (red, n = 117) to 11 other angiosperm species: *Nicotiana rustica* (n = 15 GCs), *Beta vulgaris* (n = 15), *Glycine max* (n = 21), *Solanum lycopersicum* (n = 15), *Phaseolus vulgaris* (n = 16), *Cucumis melo* (n = 17), *Lactuca sativa* (n = 13), *Nicotiana tabacum* (n = 14), *Myosotis sp*. (n = 14), *Delosperma cooperi* (n = 11), and *Leonotis leonurus* (n = 17). **(B)** Distribution of individual SW lengths in the same 11 species (n = double the number reported for each species in (A)).

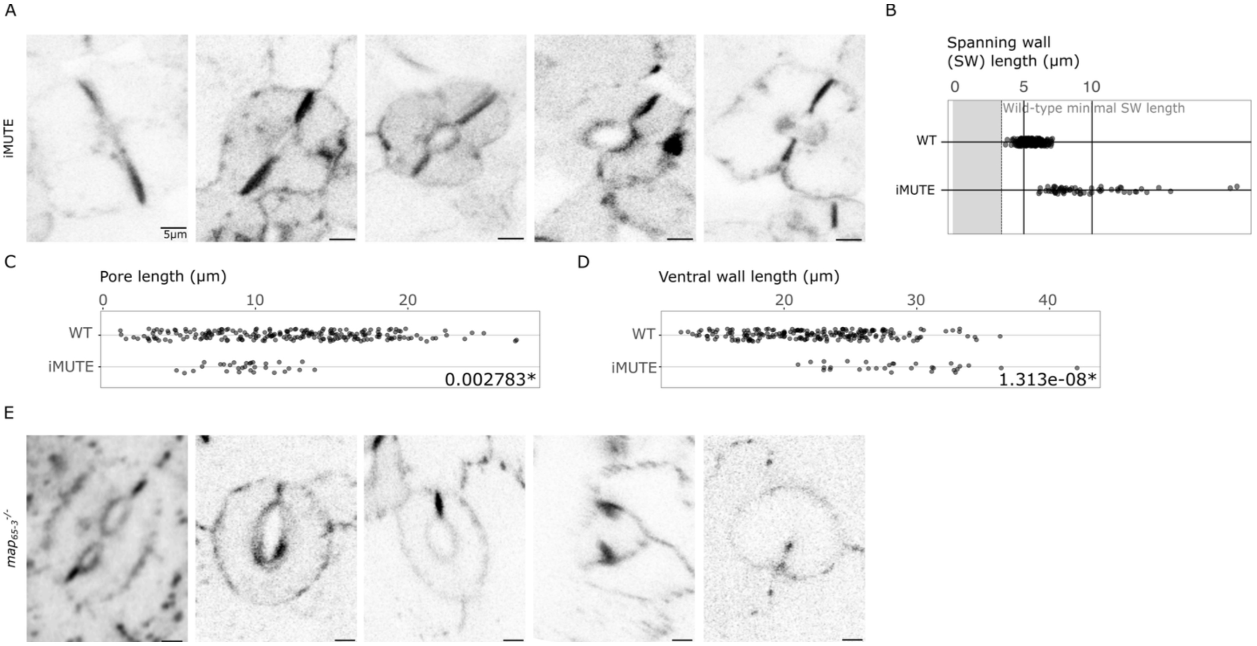


**Fig. S3 iMUTE and *map65–3* stomatal phenotypes**

**(A)** Representative aniline blue–stained (callose) images of iMUTE–induced stomata at 3 dpg. Scale bar 5 µm. **(B)** Individual SW length distributions in iMUTE–induced (n = 60) and wild type (n = 234) GCs in 3 dpg cotyledons; gray shading marks the wild type 7 dpg minimal SW length. (**C, D**) pore length (C, p.value = 0.02783*) and ventral wall length (D, p.value = 1.313e–08*) in 7 dpg cotyledons of wild type and 3 dpg cotyledons of iMUTE stomata (n = 117, and 30, respectively). **(E)** Representative aniline blue–stained images of *map65–3* stomata at 7 dpg.

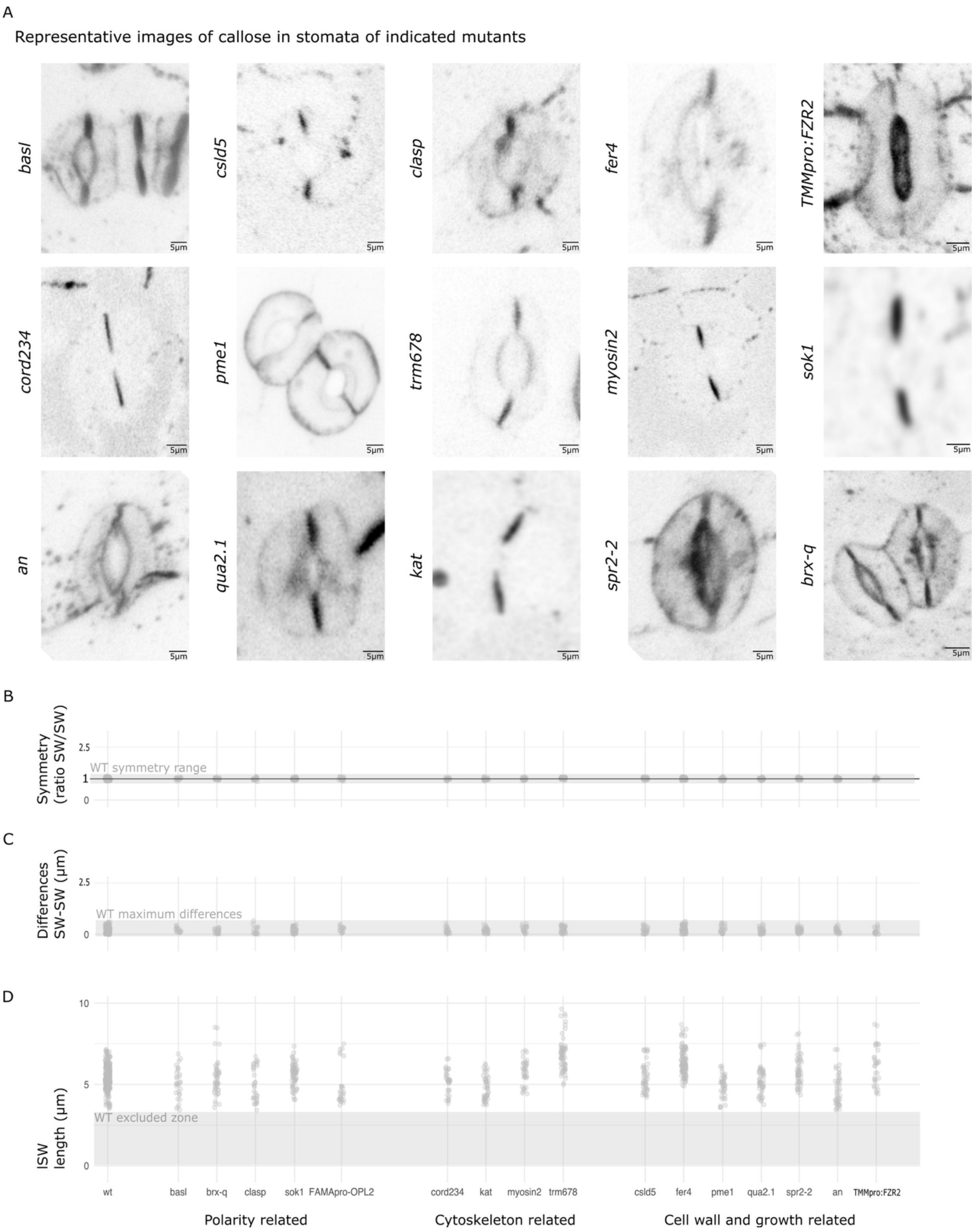
A
Representative images of callose in stomata of indicated mutants
basl
csld5
clasp
fer4
TMMpro:FZR2
cord234
pme1
trm678
myosin2
sok1
an
qua2.1
kat
spr2-2
brx-q
5µm
B
Symmetry (ratio SW/SW)
2.5
1
0
WT symmetry range
C
Differences SW-SW (µm)
2.5
0
WT maximum differences
D
ISW length (µm)
10
5
0
WT excluded zone
wt
basl
brx-q
clasp
sok1
FAMApro-OPL2
cord234
kat
myosin2
trm678
csld5
fer4
pme1
qua2.1
spr2-2
an
TMMpro:FZR2
Polarity related
Cytoskeleton related
Cell wall and growth related

**Fig. S4 Representative images and quantification of SW length and symmetry in tested mutants**

**(A)** Representative aniline blue–stained stomata from 7 dpg cotyledons of: TMMpro:FZR2 (n = 16), *clasp* (n = 16), FAMApro:OPL2 (n = 14), *trm678* (n = 31), *fer4* (n = 53), *sok1* (n = 32), *kat* (n = 23), *qua2.1* (n = 25), *pme1* (n = 20), *cord234* (n = 17), *spr2–2* (n = 27), *csld5* (n = 24), *myosin2* (n = 18), *basl* (n = 15), *angustifolia* (n = 22), *brx–q* (n = 23), *col.0* (n = 117). At least 3 independent cotyledons were imaged per genotype. **(B)** SW length ratio (d1/d2, $log_{10}$ scale) across all mutants; gray shading, maximal wild type variation. **(C)** Absolute SW length difference (|d1 − d2|) for all mutants; gray shading, maximal wild type difference. **(D)** Individual SW length distributions across all mutants; gray shading, wild type minimal SW length (WT excluded zone).

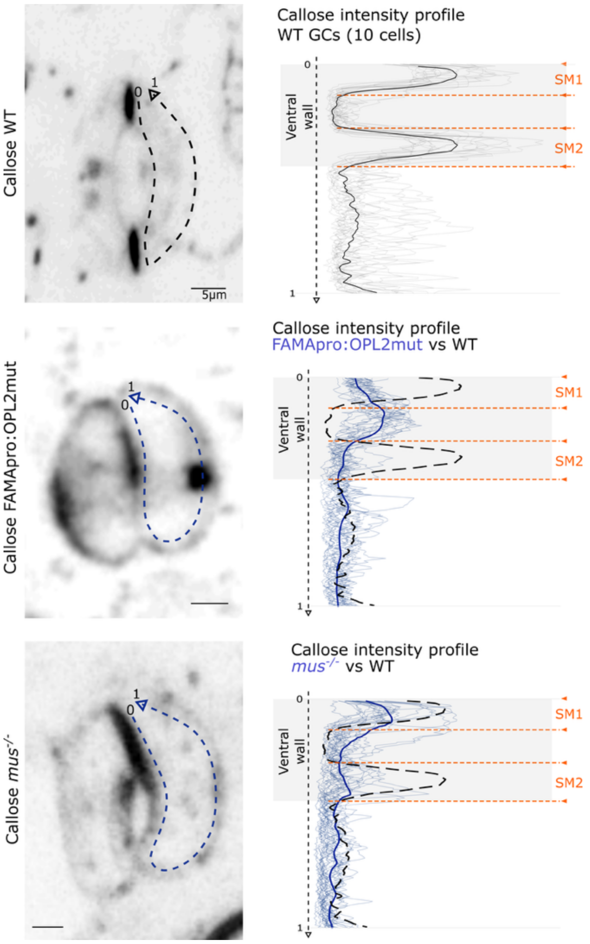


**Fig. S5 Callose intensity profiles in *mus*$^{-/-}$ mutants and OPL2 overexpression.**

Left, aniline blue–stained images of wild type, *mus*, and FAMApro:OPL2mut stomata (same images used in main figures, repeated here for ease of comparison with intensity profiles); dotted arrow indicates intensity profile orientation. Right, callose intensity profiles across the ventral wall for wild type (n = 10, mean in black) and *mus* and FAMApro:OPL2mut (n = 10 each, individual profiles in light blue, average in blue) overlaid with the WT average (black dotted profile), revealing altered callose distributions relative to wild type.

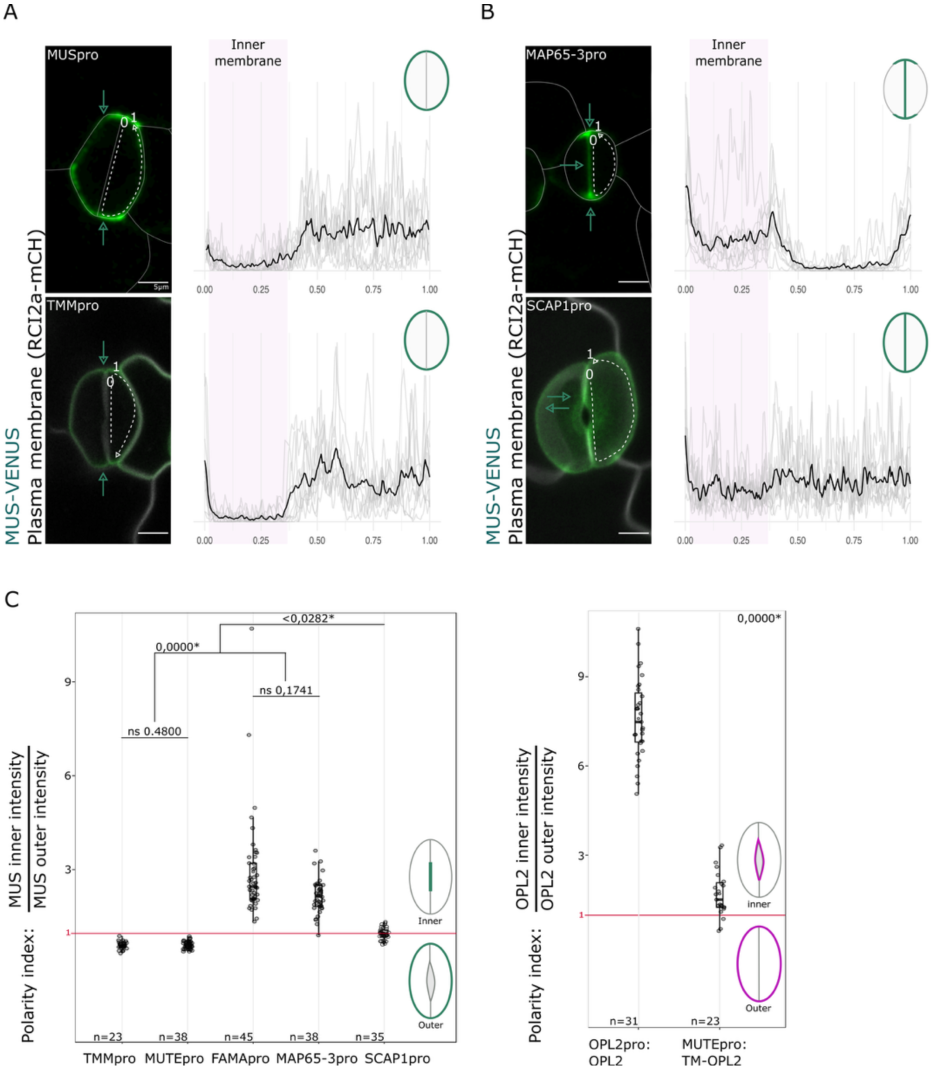


**Fig. S6 Temporal direction of MUS and OPL2 subcellular localization in GCs**

**(A)** Left, representative images of MUS–VENUS (green) driven by the early lineage promoters. The dotted white arrow indicates the orientation of the intensity profile. Right, averaged intensity profiles across the combined inner and outer membrane (n = 10 GCs, arbitrary units). Predominant GC localization of the construct cartooned on upper far right. **(B)** MUS–VENUS driven by the division specific promoter MAP65–3pro (top) or the late GC promoter SCAP1pro (bottom). **(C)** Intensity ratio of MUS, OPL2 or TM–OPL2 reporters across inner vs outer GCs membrane for each promoter used. 3 cotyledons were imaged per condition and between 23 and 45 stomata were used to extract polarity index for each line.

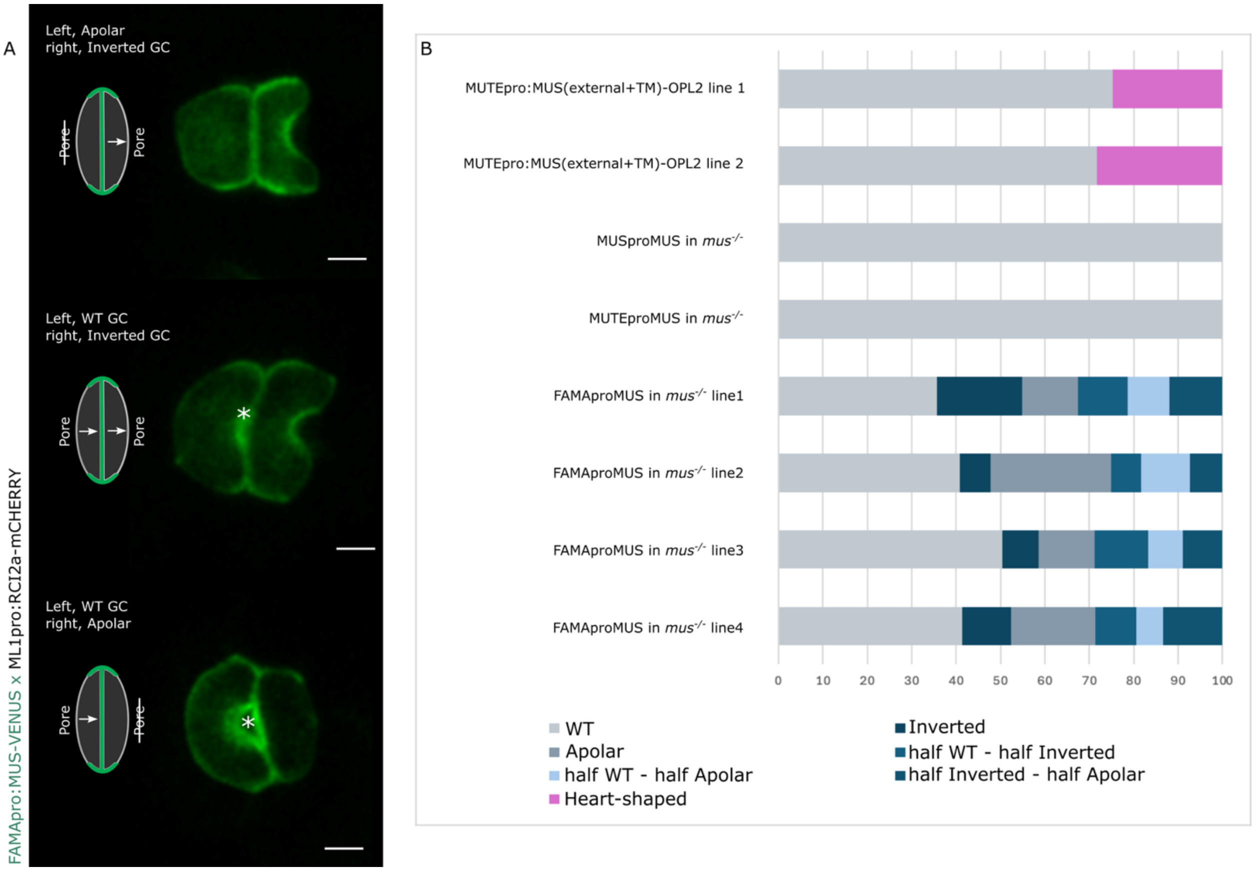


**Fig. S7 Quantification of MUS misexpression–associated defects in multiple transgenic lines.**

(**A**) Additional stomatal phenotype classes observed upon MUS misexpression, with cartoon representations of each defect class and phenotypes frequencies. (**B**) Quantification of defective stomata per transgenic line expressed as percentages. Right, all stomata defects observed are represented as percentages per line. Between 3 and 10 cotyledons were used per condition, 272 to 367 stomata counted per MUS line (in total: 130/1140 Inverted, 203/1140 Apolar, 328/1140 mixed fate). 99 and 69 stomata counted for OPL2 lines 1 and 2 (in total 45/168 Heart–shaped).

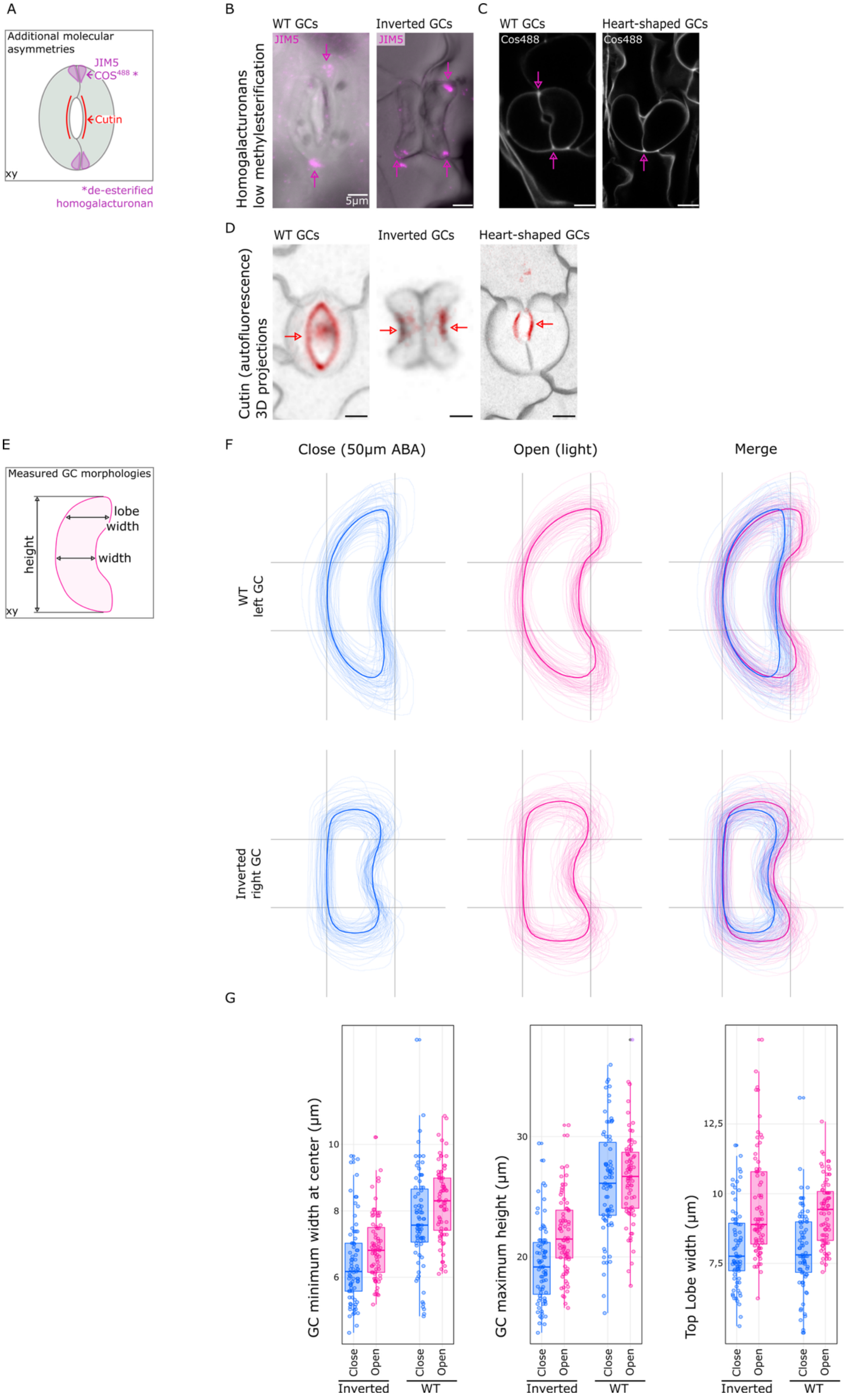

A
Additional molecular asymmetries
JIM5
COS488 *
Cutin
xy
*de-esterified homogalacturonan
B
Homogalacturonans low methylesterification
WT GCs
Inverted GCs
JIM5
5µm
C
WT GCs
Heart-shaped GCs
Cos488
D
Cutin (autofluorescence) 3D projections
WT GCs
Inverted GCs
Heart-shaped GCs
E
Measured GC morphologies
height
lobe width
width
xy
F
Close (50µm ABA)
Open (light)
Merge
WT left GC
Inverted right GC
G
GC minimum width at center (µm)
GC maximum height (µm)
Top Lobe width (µm)
Close
Open
Inverted
WT

**Fig. S8 Additional guard cell asymmetries.**

**(A)** Schematic of low–methylesterified/de–esterified homogalacturonan and cutin distribution in guard cells. **(B)** Immunolocalization of low–methyl–esterified and de–esterified homogalacturonan (magenta). Immunolocalization was consistent across all labelled wild type or inverted stomata on multiple leaves (3 WT stomata and 4 Inverted stomata were labeled). **(C)** Z–stack representative images of de–esterified homogalacturonan (COS488) staining in wild type and heart–shaped GCs. **(D)** Representative 3D projections of cutin autofluorescence (red) in wild type (RCI2a–mCH), Inverted (FAMApro:MUS–VENUS), and heart–shaped (MUTEpro:TM–OPL2, RCI2a–mCH) stomata. (3 cotyledons imaged per condition, cutin localization was consistent across all stomata observed). **(E)** Schematic of the guard cell shape parameters quantified in (F) and (G). **(F)** Extracted GC outlines from wild type (left GC) and Inverted (right GC) stomata following induced closure or opening of the pore (n = 75 GC each); mean shapes shown in blue (close) and pink (open). **(G)** Quantification of GC shape parameters (highlighted in E).

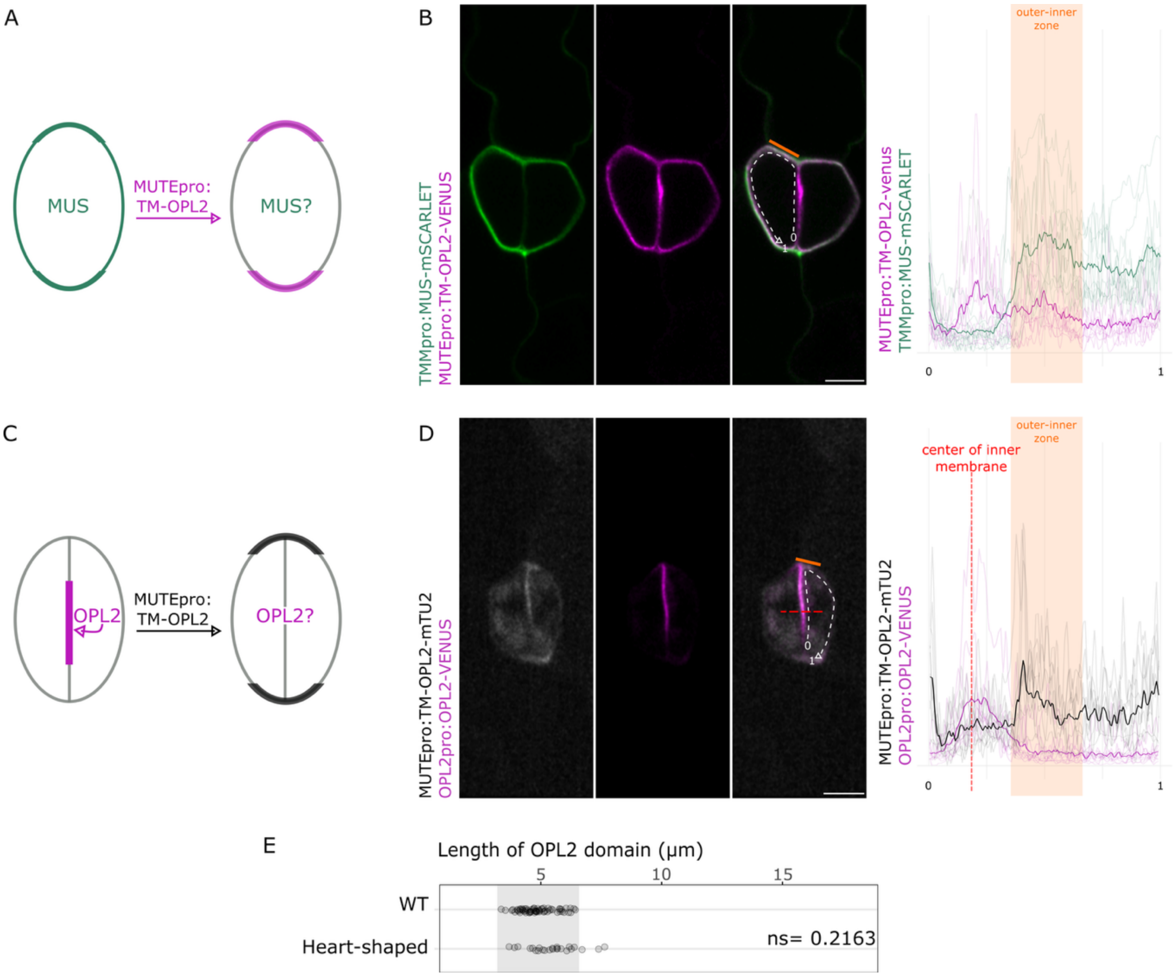


**Fig. S9 MUS localization is unaffected by OPL2 relocalization to the outer membrane.**

**(A)** Schematic of MUS localization following forced OPL2 relocalization to the outer membrane. **(B)** Representative medial section of heart–shaped stomata co–expressing MUTEpro:TM–OPL2 (magenta) and TMMpro:MUS (green), with intensity profiles comparing TM–OPL2 and MUS distributions across the inner and outer membrane. **(C)** Schematic of OPL2pro:OPL2 localization in heart–shaped stomata also expressing MUTEpro:TM–OPL2. **(D)** Representative medial section of heart–shaped stomata co–expressing MUTEpro:TM–OPL2–mTU2 (grey) and OPL2pro:OPL2–VENUS (magenta), with intensity profiles; red dotted line marks the inner membrane center to highlight the shift in OPL2 central localization. **(E)** OPL2pro:OPL2–VENUS polarity domain length in wild type (n = 64 GCs) and heart–shaped (n = 24, 3 cotyledons) stomata.

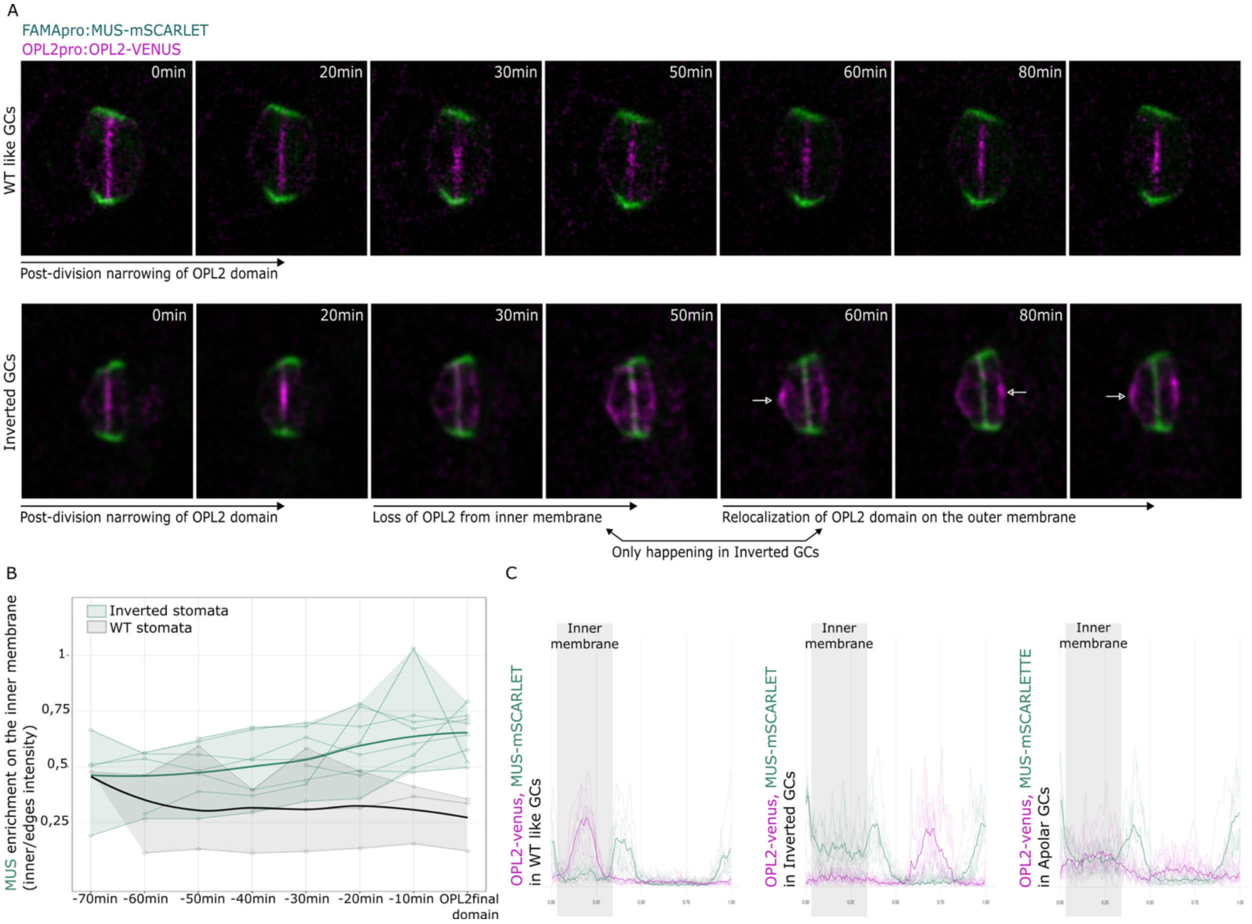


**Fig. S10 Timelapse tracking of the reorganization of GC polarity upon MUS mislocalization**

**(A)** Timelapses of OPL2pro:OPL2 (magenta) localization upon expression of FAMApro:MUS (green). Normal stomata (top) result when OPL2 remains in the inner membrane domain and is not replaced by MUS. Inverted stomata (bottom row) form when MUS invades the inner membrane and OPL2 is displaced on the outer membrane, as marked by white arrows. **(B)** Quantification of MUS relocalization (inner membrane intensity/edge intensity) during polarity establishment, until OPL2 reaches its final localization on the inner (wild type–like, n = 3 cells) or outer (Inverted, n = 9 cells) membrane. **(C)** OPL2 and MUS intensity profiles across GC membrane in wild type–like (left), Inverted (middle), and Apolar (right) guard cells (n = 10 cells each).

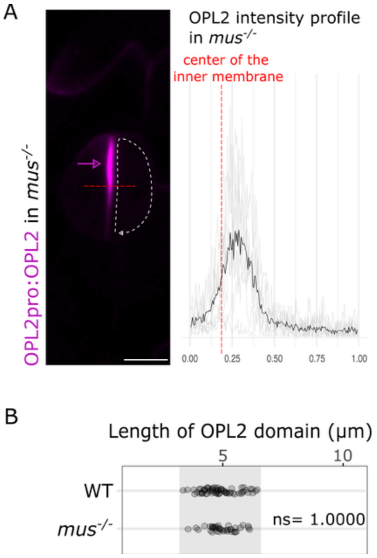


**Fig. S11 OPL2 generates a gradient of consistent length and peak dimensions in the absence of MUS.**

**(A)** Representative medial section of *mus*$^{-/-}$ stomata expressing OPL2pro:OPL2–VENUS with corresponding intensity profile; red line marks the inner membrane center. **(B)** OPL2pro:OPL2–VENUS polarity domain length in wild type (n = 64) and *mus*$^{-/-}$ (n = 38 stomata, from 5 cotyledons).

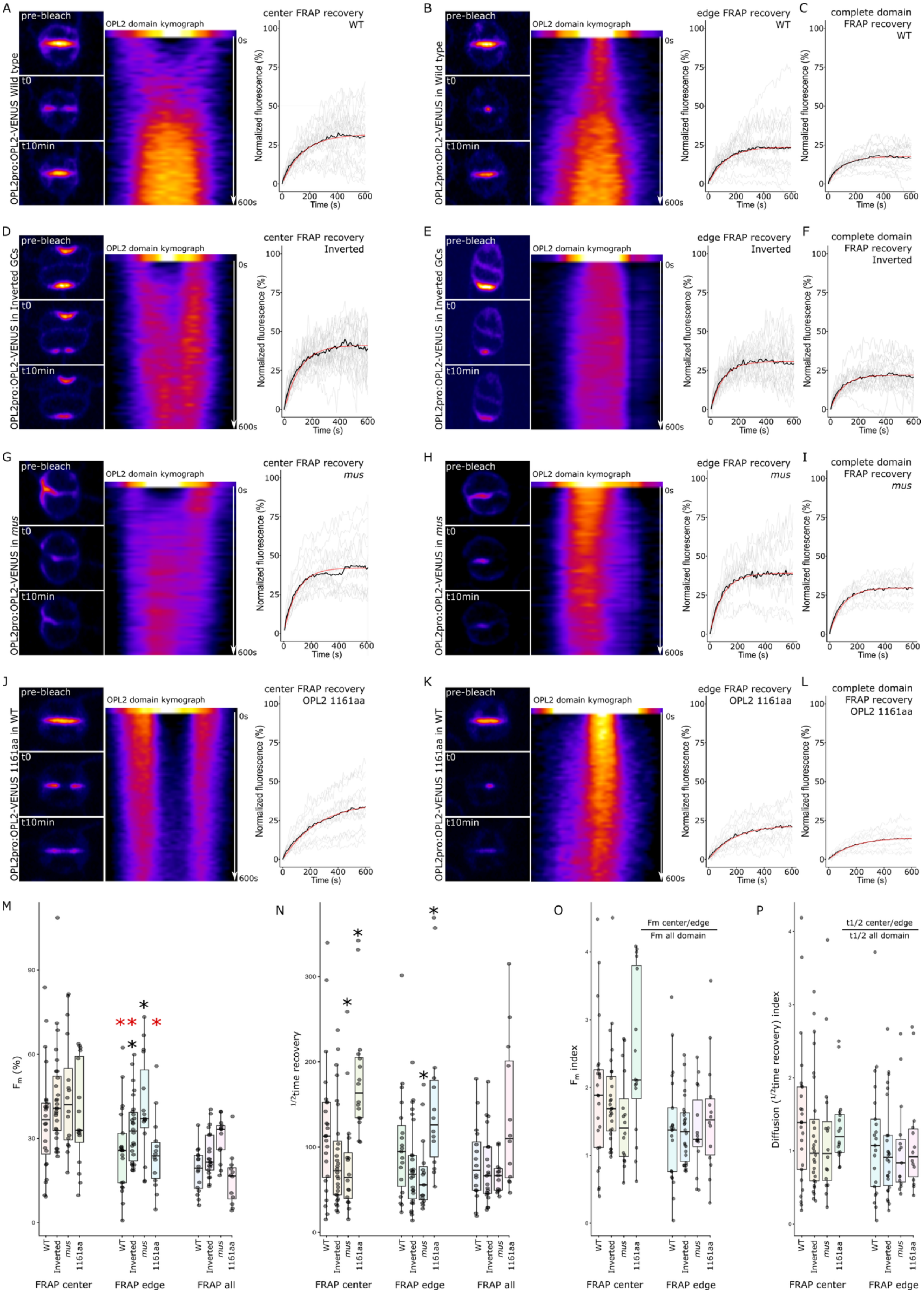
A
pre-bleach
t0
t10min
OPL2 domain kymograph
0s
600s
OPL2pro:OPL2-VENUS Wild type
center FRAP recovery
WT
Normalized fluorescence (%)
Time (s)
B
OPL2pro:OPL2-VENUS in Wild type
edge FRAP recovery
WT
C
complete domain
FRAP recovery
WT
D
OPL2pro:OPL2-VENUS in Inverted GCs
center FRAP recovery
Inverted
E
edge FRAP recovery
Inverted
F
complete domain
FRAP recovery
Inverted
G
OPL2pro:OPL2-VENUS in mus
center FRAP recovery
mus
H
edge FRAP recovery
mus
I
complete domain
FRAP recovery
mus
J
OPL2pro:OPL2-VENUS 1161aa in WT
center FRAP recovery
OPL2 1161aa
K
edge FRAP recovery
OPL2 1161aa
L
complete domain
FRAP recovery
OPL2 1161aa
M
Fm (%)
WT
Inverted
mus
1161aa
FRAP center
FRAP edge
FRAP all
N
1/2time recovery
O
Fm center/edge
Fm all domain
Fm index
P
t1/2 center/edge
t1/2 all domain
Diffusion (1/2time recovery) index

**Fig. S12 FRAP analysis reveals that OPL2 domain dynamics are influenced by MUS.**

All panels show FRAP recovery analysis of OPL2pro:OPL2–VENUS in GCs of true leaves. Unless otherwise noted, each panel shows three representative timelapse images (pre–bleach, t = 0, t = 10 min), a kymograph of inner membrane intensity over 10 min (pre–bleach duplicated for visualization), and individual recovery curves (grey) with the average in black and the fitted curve in red; panels with complete–domain ROIs show recovery curves only (**C, F, I, L**). **(A)** Wild type, center ROI (n = 24). **(B)** Wild type, edge ROI (n = 21). **(C)** Wild type, complete domain FRAP (n = 16). **(D)** Inverted, center ROI (n = 28). **(E)** Inverted, edge ROI (n = 26). **(F)** Inverted, complete domain FRAP (n = 21). **(G)** *mus*, center ROI (n = 16). **(H)** *mus*, edge ROI (n = 15). **(I)** *mus*, complete domain FRAP (n = 13). **(J)** Extended length (300 extra aa for total of 1161 aa) OPL2pro:OPL2–VENUS, center ROI (n = 16). **(K)** OPL2pro:OPL2–VENUS (1161 aa), edge ROI (n = 14). **(L)** OPL2pro:OPL2–VENUS (1161 aa), complete domain FRAP (n = 12). **(M)** Mobile fraction (Fm) at 10 min post–bleach for all conditions and ROI positions. **(N)** Half–recovery time (t½) from curve fitting for all conditions. **(O)** Fm index: Fm of center or edge ROI divided by average Fm of entire–domain FRAP, highlighting relative contributions of lateral diffusion vs. cytosolic exchange. **(P)** Diffusion index: t½ of center or edge ROI divided by average t½ of entire–domain FRAP. Black asterisks, significant differences vs. wild type; red asterisks, significant differences between center and edge within the same condition.

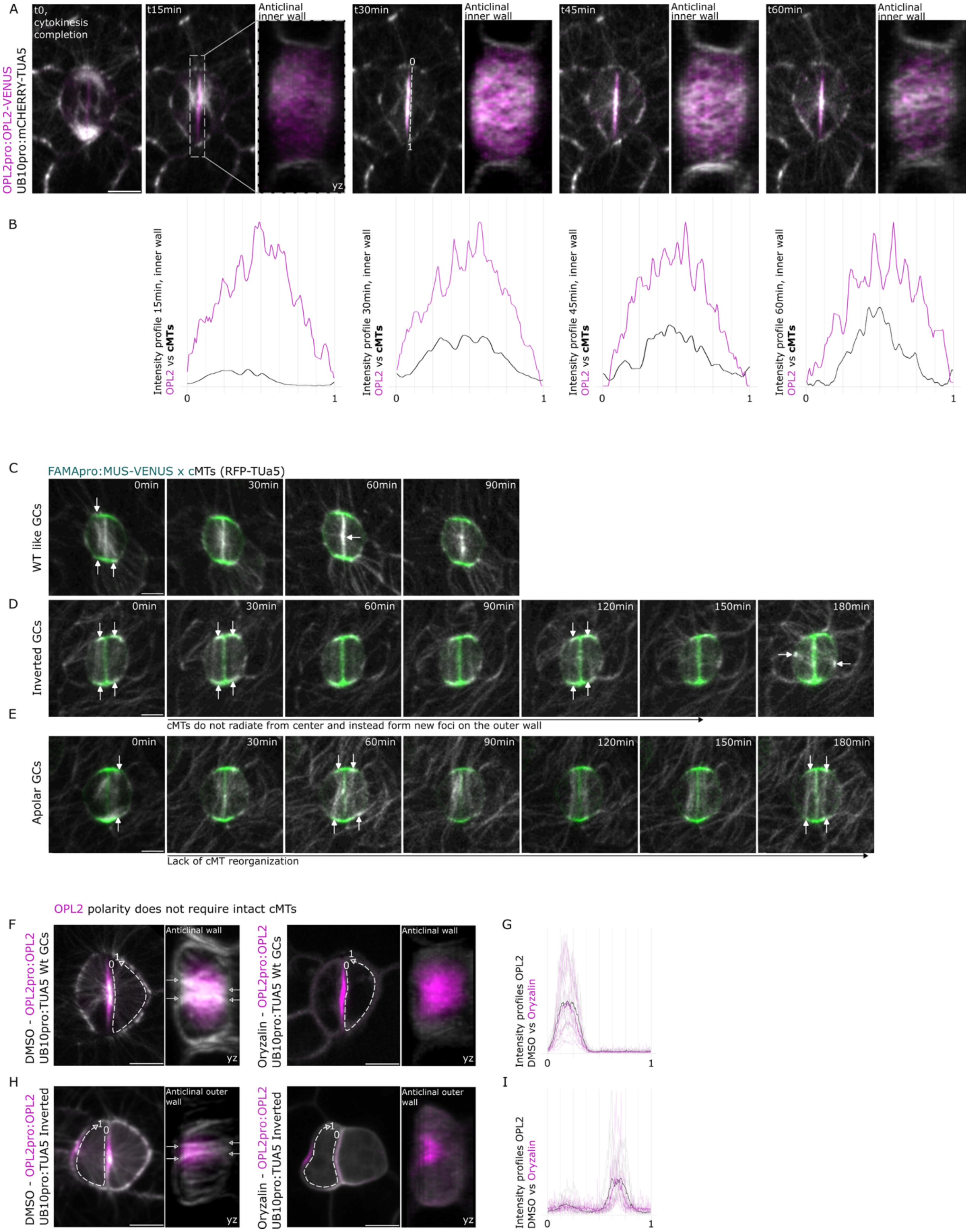
A
OPL2pro:OPL2-VENUS
UB10pro:mCHERRY-TUA5
t0, cytokinesis completion
t15min
Anticlinal inner wall
yz
t30min
t45min
t60min
B
Intensity profile 15min, inner wall
OPL2 vs cMTs
Intensity profile 30min, inner wall
Intensity profile 45min, inner wall
Intensity profile 60min, inner wall
C
FAMApro:MUS-VENUS x cMTs (RFP-TUa5)
WT like GCs
0min
30min
60min
90min
D
Inverted GCs
120min
150min
180min
cMTs do not radiate from center and instead form new foci on the outer wall
E
Apolar GCs
Lack of cMT reorganization
OPL2 polarity does not require intact cMTs
F
DMSO - OPL2pro:OPL2
UB10pro:TUA5 Wt GCs
Anticlinal wall
Oryzalin - OPL2pro:OPL2
G
Intensity profiles OPL2
DMSO vs Oryzalin
H
UB10pro:TUA5 Inverted
Anticlinal outer wall
I

**Fig. S13 Dynamic coupling between microtubules and guard cell polarity.**

**(A, B)** Timelapse imaging of cMTs (grey) and OPL2pro:OPL2 (magenta) following guard cell division in wild type true leaves (15 min intervals). (**A**) Maximum projection (left: xy view; right: yz view). (**B**) Associated intensity profiles of cMTs (black) and OPL2 (magenta) across the inner membrane. **(C–E)** Timelapse imaging of cMTs (grey) with FAMApro:MUS (green), shown for wild type–like GCs (**C**), Inverted GCs (**D**), and Apolar GCs (**E**). (**F, H**) Maximum projections of OPL2pro:OPL2–VENUS (magenta) and UB10pro:mCHERRY–TUA5 (grey; left: xy view; right: yz view) in 3 dpg wild type (**F**) and Inverted GCs (**H**) treated with DMSO (left) or 20 μM Oryzalin for 2 h (right). (**G, I**) Corresponding OPL2 intensity profiles across the GC membrane (DMSO, grey; Oryzalin, magenta) for every condition listed previously.

**Criteria**

*1- Stereotyped domain size*
*2- Central OPL2 peak*
*3- >10× OPL2 → pole-to-pole spread*
*4- OPL2 self-organizes without MUS*
*5- MUS clears established OPL2*
*6- OPL2 can fail to repolarize*
*7- OPL2 does not relocalize MUS*
*8- TM-OPL2 bypasses MUS control*
*9- MUS and OPL2 remain separated*
*10- No puncta; displaced OPL2 is diffuse*
*11- Center Fm = edge Fm in mus$^{-/-}$*
*12- Whole-domain Fm differs between conditions*
*13- Center Fm > edge Fm in WT and Inverted*
*14- No center/edge t½ difference per genotype*

| Model/schematic representation | Fit score |
|---|---|
| **A-Physical confinement / "picket-fence" corral at the edge** | |

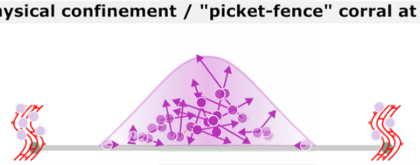

10.5/14: 2 contradictions

✗ Disqualifying prediction: direct contact between OPL2 and MUS is required (*criteria 9*).

**B- Boundary-localized capture-and-clearance**

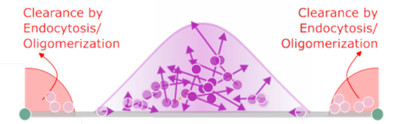


10/14: 2 contradictions

✗ Disqualifying prediction: accumulation of immobile OPL2 puncta on edges is required (*criteria 11*).

**C- Polarized delivery and an independent edge trap**

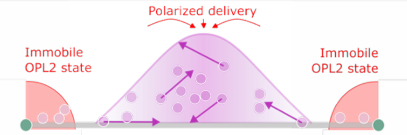


11/14: 6 partial fit

~ Untested requirement: delivery dependency on MUS with either convergent independent pathways or a dual MUS role (centering + edge-trapping) .

**D- Advective/directed transport toward the center**

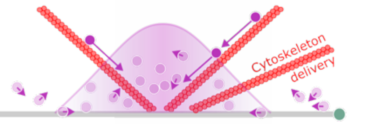


7/14: 5 contradictions

✗ Disqualifying prediction: requires a center–edge t½ difference (*criteria 14*) + transport independent of OPL2 size (*criteria 13*).

**E- Emergent front dynamics in a bistable, mass-conserved system**

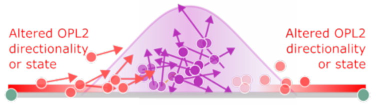


13/14: 2 partial fit

~ Untested requirement: MUS alteration of OPL2 state implies (re)directing OPL2 toward the center.

**F- Direct binding/tether**

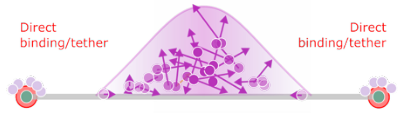


8.5/14: 4 contradictions

✗ Disqualifying prediction: direct contact between OPL2 and MUS, where OPL2 binds to MUS, is required (*criteria 1, 5-6, 9*).

**G- Steric exclusion**

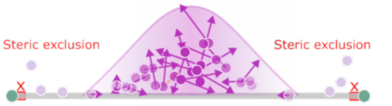


3/14: 10 contradictions

✗ Disqualifying prediction: direct contact between OPL2 and MUS is required, colocalization is not possible (*criteria 3-6, 8-14*).

**H- Complete independency from MUS**

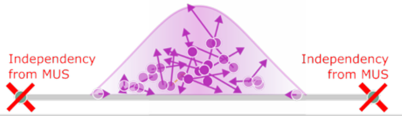


6/14: 7 contradictions

✗ Disqualifying prediction: OPL2 should be insensitive to MUS (*criteria 5, 8-14*).

OPL2 mobility → OPL2 activity state + ●●●● - OPL2 density max MUS localization ● MUS function max

**Fig. S14 Comparative evaluation of eight candidate mechanistic models for MUS–OPL2 domain lengths and behaviors**

Eight candidate mechanistic models were scored against 14 discrete experimental observations or data–derived predictions (top) drawn from live–cell reporter imaging, genetic, and FRAP data. Left, schematic representation of each model, with the proposed MUS–dependent behavior highlighted in red. Right, tallied fit score and model–disqualifying prediction. Model E best fits for current data. Additional explanation of the models is provided in methods.

### Movie S1– Timelapse of cMTs and OPL2.

Time–course analysis of cMTs and OPL2 rearrangement from GMC to pore formation. (10 min intervals)

### Movie S2– 3D projection of Inverted GCs.

3D rotation of Inverted GCs expressing MUS–mSCARLET, mCHERRY–TUa5 and OPL2–VENUS.

### Movie S3– Timelapse of cMTs, FAMApro:MUS and OPL2.

Time–course analysis (20 min intervals) of cMTs rearrangement (3D projection) following OPL2 establishment on the outer membrane (right GC of both stomata).
